# Bacterial extracellular vesicles indirectly destabilize a human stem cell–derived blood–brain barrier on-chip through pro-inflammatory stimulation of immune cells

**DOI:** 10.64898/2026.01.23.701361

**Authors:** Louis P. Widom, Panteha Torabian, Michelle A. Trempel, Molly C. McCloskey, Lea V. Michel, James L. McGrath, Thomas R. Gaborski

## Abstract

Pathogenic bacterial extracellular vesicles (BEVs) can disrupt the blood–brain barrier (BBB), leading to neuroinflammation. Prior *in vitro* studies of this process were performed in simple models that may have lacked important physiological factors. We sought to determine if treatment with *Escherichia coli*–derived BEVs could directly compromise the integrity of a BBB lab-on-chip model or if an immune component was required. Our device featured isogenic human induced pluripotent stem cell–derived brain microvascular endothelial-like cells (BMECs) and pericytes separated by an ultrathin, porous silicon nitride membrane. BEVs and free lipopolysaccharide (LPS) were capable of causing upregulation of intercellular adhesion molecule-1 on the BMEC surfaces, which is important for immune cell recruitment. However, neither BEVs nor LPS at physiological doses caused pronounced loss of BMEC tight junction proteins, nor did they increase barrier permeability to small dye molecules. In contrast, stimulating THP-1 macrophages with BEVs led to increased production of pro-inflammatory cytokines, and conditioned media from the stimulated macrophages disrupted BMEC tight junctions and increased barrier permeability. Our work demonstrates the importance of incorporating an immune component in studies of BEV-mediated disruption of BBB models.

## Introduction

Sepsis is a potentially fatal organ disruption stemming from a hyperactive inflammatory response to infection,^1^ and it is a leading cause of mortality in hospitals.^2^ The invasion of bacteria or other pathogens causes the immune system to increase production of pro-inflammatory signals, leading to a cytokine storm that can initiate fever and cause host cell death.^3,4^ This dysregulated host response can be more harmful than the invading pathogens due to self-injury of tissues and organs.^1,4^ Additionally, sepsis can cause widespread endothelial dysfunction, which is especially dangerous in the brain vasculature.^5^

The interface between the vasculature and brain parenchyma is termed the blood–brain barrier (BBB) and it defends the central nervous system by exerting size, chemical, and charge-based selection to prevent entry of disruptive molecules and cells while still allowing the exchange of oxygen and essential nutrients.^6^ Brain microvascular endothelial cells are primarily responsible for this neuroprotective role. Neighboring endothelial cells form tight junctions composed of zonula occludens-1, claudin-5, and occludin that restrict paracellular diffusion across the BBB.^6^ Furthermore, brain microvascular endothelial cells possess efflux proteins that are embedded in the cell plasma membranes and prevent transcellular transport of lipophilic molecules by pumping them back into the blood.^7^

Other cell types support the specialized function of brain microvascular endothelial cells. Pericytes are one of the mural cells that surround the vasculature. Their contractile nature helps regulate blood flow,^8^ and their communication with endothelial cells may strengthen barrier properties.^9–13^ Pericytes are embedded within the vascular basement membrane,^14^ a collection of extracellular matrix proteins that surrounds the basolateral surface of blood vessels and supports the endothelium. The basement membrane is composed of collagen IV and laminin networks joined together by smaller proteins.^15^ Production of the basement membrane is enhanced by the presence of pericytes,^13,16–19^ and since evidence suggests that endothelial interactions with the basement membrane strengthen expression of tight junction proteins,^20^ this may be one way in which pericytes support the BBB.

During sepsis, pro-inflammatory cytokines cause breakdown of endothelial tight junctions and promote barrier leakiness.^21,22^ They also upregulate intercellular adhesion molecule-1 (ICAM-1) on the surface of endothelial cells.^23^ This 80–114 kDa transmembrane protein^24^ is likewise upregulated on leukocytes such as macrophages under pro-inflammatory conditions, where it is believed to play a role in phagocytosis.^25^ In the context of endothelial ICAM-1, this protein serves to recruit leukocytes from the circulation.^23^ Once leukocytes bind to the endothelium, they can transmigrate into the brain to promote neuroinflammation by activating brain-resident microglia.^26,27^ Pericytes may help resist invasion by leukocytes,^18^ but the increased pro-inflammatory signaling during sepsis can cause pericyte loss from the BBB which would reduce this neuroprotective effect.^28,29^ Neuroinflammation that arises during sepsis can result in delirium.^27^ A major concern for patients is that the BBB can remain disrupted following clearance of sepsis, and sepsis survivors consequently have a higher risk of developing neurological disorders.^5^

Bacterial extracellular vesicles (BEVs) are bacterial lipid membrane–derived nanoparticles that contribute to sepsis pathogenesis.^30^ Their cargo includes signaling factors, nucleic acids, lipopolysaccharide (LPS), and other toxins incorporated from the source bacteria.^31–33^ As such, exposure to these 40–400 nm^34^ particles can upregulate ICAM-1 in endothelial cells^35–37^ and increase pro-inflammatory cytokine production by leukocytes.^38,39^ BEVs may compromise BBB integrity by disrupting endothelial tight junctions and increasing paracellular permeability, which has been observed both *in vivo*^40,41^ and *in vitro*.^42,43^ This is one proposed mechanism that could allow BEVs to enter the brain, where they could exacerbate neuroinflammation.^44^ Prior work has been useful for examining the direct impact of pathogenic BEVs on *in vitro* human BBB models, but, to our knowledge, these direct impacts have not yet been compared to indirect impacts on BBB integrity mediated by BEV interactions with leukocytes. We sought to explore this gap using a lab-on-chip model of the human BBB.

Many prior *in vitro* studies of the BBB have involved culturing brain endothelial cells on porous membranes to model the blood vessel wall.^11,13,20,45,46^ Recent advances in the manufacturing of porous membranes has led to development of the microphysiological system enabled by an ultrathin silicon membrane (µSiM) barrier modeling platform.^47–49^ This modular lab-on-chip device incorporates a continuously nanoporous silicon nitride membrane that is ∼100 nm thick, similar to the physiological vascular basement membrane and orders of magnitude thinner than traditional culture inserts.^15,48^ As a result, cells seeded on either side of the membrane are placed in close proximity, reducing the distance required for the exchange of signaling factors and improving cell communication and barrier integrity.^50^ The other benefit of this technology is the optical transparency of the silicon nitride, which facilitates imaging directly in the µSiM device. Several published studies have demonstrated the µSiM’s utility for modeling the BBB.^18,19,49,51–53^

In the present work, we used isogenic extended endothelial culture method brain microvascular endothelial cell-like cells (BMECs)^54^ and brain pericyte-like cells (BPLCs)^55^ derived from the same parent human induced pluripotent stem cell (hiPSC) line. Together, these cell co-cultures formed a BMEC-pericyte barrier (BPB) that modeled the primary BBB at the post-capillary venule, which serves as the inflammatory nexus of the brain microvasculature.^56,57^ While previous investigations with the µSiM examined barrier disruption caused by exposure to pro-inflammatory cytokines^18,19,48,53^ and LPS,^58^ this is the first time the µSiM has been used to study the effects of BEVs on a barrier model. Many models of BEV stimulation have been created *in vivo*.^44^ However, animal models do not always predict human responses, motivating *in vitro* approaches using human cells.^59–62^ Our work is one of the first to incorporate both pathogenic BEVs and immune cell secretions in a lab-on-chip human BBB model. We found that BMECs were sensitive to BEVs. However, direct stimulation with BEVs failed to disrupt tight junction proteins or permeability, contradicting previous studies.^42,43^ The barrier was compromised only when the model was exposed to secretions from macrophages that had been treated with BEVs. These results suggest that including immune components is essential for *in vitro* studies to accurately predict the effects of BEV treatment on human BBB models.

## Materials and Methods

### µSiM assembly

Modular µSiM (**micro**physiological system enabled by a **si**licon **m**embrane) devices were assembled as previously described.^49^ Briefly, nanoporous silicon nitride chips with ∼60 nm diameter pores, 15% porosity, and 100 nm membrane thicknesses (SiMPore, NPSN100-1L) were used as the basis for the model. Membranes were bonded to a custom acrylic component (ALine, Inc., Signal Hill, California, USA) with pressure sensitive adhesive, and additional pressure sensitive adhesive was used to attach an underlying microfluidic channel manufactured from cyclic olefin. µSiM devices were sterilized under UV for at least 20 min following assembly, then were flipped and exposed to UV for an additional 20 min. 150 mm petri dishes were used to hold the devices, and overturned caps from 50 ml centrifuge tubes were placed in the petri dishes to serve as water reservoirs to help maintain local humidity.

### Cell culture

All cells were maintained in a 37 °C environment with 5% CO_2_.

BMECs were differentiated from IMR90-4 hiPSCs (WiCell, Madison, Wisconsin, USA) following the method developed by Nishihara *et al*.^54^ Briefly, the hiPSCs were differentiated into endothelial progenitor cells over 3 days after seeding at approximately 30,000 cells cm^-2^ in mTeSR1 media (STEMCELL Technologies, 85850). Media was then switched to Advanced DMEM/F12 (Gibco, 12634010) with GlutaMAX Supplement (Gibco, 35050061) and L-ascorbic acid for 5 days with the addition of 8 µM CHIR99021 (Sigma-Aldrich, SML1046) during the first 2 days. CD31^+^ cells were isolated via magnetic-activated cell sorting and were seeded in 6-well plates coated with 100 µg type IV collagen (Sigma-Aldrich, C5533-5MG) per well. These cells were maintained in hECSR medium, which consisted of Human Endothelial-SFM (Gibco, 11111044) supplemented with 1X B-27 Supplement (Gibco, 17504044) and 20 ng ml^-1^ human fibroblast growth factor 2 (R&D Systems, 233-FB). Media was exchanged every 1–2 days. BMECs were obtained through selective passaging up to passage 3 and used for experiments up to passage 7.

BPLCs were derived from the same progenitor hiPSCs as the BMECs and were therefore isogenic to the endothelial cells. The differentiation method was developed by Gastfriend *et al*.^55^ Briefly, the hiPSCs were differentiated into neural crest stem cells through maintenance in E6 medium (Gibco, A1516401) + CHIR99021, SB431542 (Tocris, 1614), human fibroblast growth factor 2, dorsomorphin (Sigma-Aldrich, P5499), and heparin (Sigma-Aldrich) for 15 days. p75-NGFR^+^ cells were isolated via magnetic-activated cell sorting and plated with E6 medium + 10% fetal bovine serum (FBS) (Peak Serum, PS-FB1) for the remaining culture period, and were considered BPLCs between days 22–45 of culture. Media was exchanged every 1–2 days.

Human leukemia monocytic type 1 (THP-1) cells (ATCC) were obtained from the lab of Dr. Karin Wuertz-Kozak. Cells were maintained in suspension in RPMI 1640 (Cytiva, SH30255.01) supplemented with 10% FBS (Cytiva, SH30070.03), 1% anti-anti (Gibco, SV30079.01), and 1% sodium pyruvate (Gibco, 11360-070). Media was exchanged every 2–3 days. For differentiation into M0 macrophages, the THP-1 media was supplemented with 100 ng ml^-1^ phorbol 24-myristate 13-acetate (Cayman Chemical, 16561-29-8) for approximately 48 h prior to experimentation.

### Seeding µSiM co-culture

Each µSiM device was washed twice with sterile water (100 µl in the top well, 20 µl in the lower channel) prior to the addition of coating solutions. 20 µl of 800 µg ml^-1^ collagen IV was used to coat the lower channel while the top well was coated with 100 µl of 400 µg ml^-1^ collagen IV + 100 µg ml^-1^ fibronectin (Sigma-Aldrich, F1141-1MG) at 37 °C for at least 30 min before cell seeding.

The top well and lower channel were washed with E6 medium + 10% FBS before adding BPLCs to the lower channel at a density of 14,000 cells cm^-2^. Devices were then inverted for at least 2 h to allow the BPLCs to adhere to the underside of the chip. Media remains in the top well during the inversion step due to surface tension. Following this incubation period, devices were turned upright and the top well and lower channel were washed with E6 medium + 10% FBS to remove any dead or unattached cells. 1 day later, the media in the top well and lower channel was replaced with hECSR medium, and BMECs were seeded in the top well at a density of 40,000–50,000 cells cm^-2^. After at least 2 h, media in the top well was exchanged to remove any dead or unattached BMECs. hECSR medium was exchanged every day until the cells were ready for experimentation, which was 6–7 days after BMEC seeding.

### BEV Isolation

*Escherichia coli* (*E. coli*) from strain CFT073 [WAM2267] were cultured overnight on Lysogeny Broth (LB) agar at 37 °C. Colonies were taken from this plate and added to 50 ml LB and cultured overnight at 37 °C, shaking at 160 rpm. The overnight culture was used to inoculate two larger 200 ml cultures. After their optical density reached ∼0.8 at 600 nm, the two 200 ml cultures were combined before splitting into 8 sterile flasks, which were incubated at 37 °C for another 3 hours, shaking at 180 rpm. They were then centrifuged at 5,000×g for 15 min to pellet the *E. coli*. The supernatant was passed through a 0.22 µm syringe filter and 30 ml of the filtrate was then ultracentrifuged at 100,000×g for 1 h. After removing the supernatant, BEV pellets were resuspended in 400 µl PBS so that they were brought to 75 times their original LB concentration. To confirm that no *E. coli* were co-isolated with the BEVs, aliquots of the BEV preparations were incubated on LB agar overnight, and no visible colonies formed.

### Nanoparticle tracking analysis (NTA)

Nanoparticle sizes and concentrations were measured using a NanoSight NS300 (Malvern Panalytical, Malvern, UK) equipped with a sCMOS camera, 532 nm green laser, and a 565 nm long pass filter. For each sample, three 30 s videos were recorded in clear scattering mode with a camera level of 15 and a detection threshold of 5. The instrument software calculated particle diameters based on the diffusion coefficients determined from the particle tracks in the videos. Samples were further diluted in PBS to ensure that they fell within the instrument’s accurate concentration detection range of 1×10^8^–1×10^9^ particles ml^-1^. The average particle concentrations reported in the present study take the dilution factors into account.

### Transmission electron microscopy (TEM)

A PELCO Easiglow was used to discharge 200 mesh copper grids coated with formvar/carbon film for 30 s at 30 mA before adding 3 μl of the liquid BEV sample for 30 s. After wicking away excess sample, the grids were washed three times with 15 μl molecular grade water. Negative staining was performed with two applications of 10 μl filtered 0.75% uranyl formate and excess fluid was wicked away using hardened Whatman 50 filter paper between steps. Following a drying step, the grids were examined on a Talos 120C transmission electron microscope with a CETA 16 megapixel camera (Thermo Fisher) for image capture using TIA (Thermo Fisher).

### Western blot

To obtain *E. coli* whole cell lysates, *E. coli* were cultured on LB agar at 37 °C overnight. Cells were transferred to a microtube containing 50% PBS + 50% RIPA lysis buffer. BEVs were also transferred to a 50% PBS + 50% RIPA solution. Both suspensions were sonicated for 2 min. The tubes were then spun down at 5000×g for 5 min to pellet larger debris, and the supernatants were transferred to fresh tubes and stored at -20 °C until ready. The Micro BCA Protein Assay Kit (Thermo Fisher, 23235) was used to determine protein concentrations in the lysate samples. Following the manufacturer’s protocol, bovine serum albumin standards and the samples were diluted in 50% PBS + 50% RIPA buffer and loaded into a multiwell plate along with the assay working reagent. Following 2 h incubation at 37 °C, the plate’s absorbance was measured at 562 nm. After subtracting the absorbance value measured from blank wells, a polynomial fit was applied to the standards to generate a curve from which the sample protein concentrations were calculated.

The concentrations obtained from BCA were used to determine the volumes required to run 1 µg protein per sample for western blot. Samples were mixed with Laemmli buffer + 10% β-mercaptoethanol and heated at 95 °C for 5 min. After loading the samples in a 4–20% Mini-PROTEAN® TGX™ Precast Protein Gel (Bio-Rad, 4561093), 150V was applied for 30 min. Proteins were transferred to a 0.2 µm nitrocellulose membrane using a Trans-Blot® Turbo™ Transfer System (Bio-Rad, 1704150). 5% BSA mixed in tris-buffered saline (TBS) was used to block the membrane for 1 h before three subsequent washes with TBS. To label peptidoglycan-associated lipoprotein (Pal), the membrane was incubated with anti-Pal antisera from mice inoculated with purified recombinant non-lipidated Pal (∼21 kDa; contains a 6xHis-tag for purification; Rochester General Hospital)^63^ diluted 1:7,000 in TBS + 0.1% Tween-20 (TBS-T) on a shaker overnight at 4 °C. After three washes with TBS-T, the membrane was incubated with IRDye® 800CW Goat anti-Mouse IgG1-Specific Secondary Antibody (LICORbio, 926-32350) diluted 1:20,000 in 5% BSA in TBS-T on a shaker for 1 h at room temperature. The membrane was washed 3 more times with TBS-T and then imaged using a LICOR Odyssey XF Imager with a 30 s exposure in the 800 nm channel.

### Endotoxin quantification

The Pierce™ Chromogenic Endotoxin Quant Kit (Thermo Fisher, A39552) was used to measure the amount of LPS in our BEVs. Briefly, the kit’s endotoxin standard was vortexed for 15 min in water. A series of comparison standards was generated by serial dilution. Endotoxin-free water was also used to serially dilute the BEV sample. After pre-heating a multiwell plate for 10 min at 37 °C, we added 50 µl of the standards and samples. Amebocyte lysate reagent was then reconstituted in water and 50 µl was added to the wells. The solutions were mixed by tapping the plate, and the reaction ran for 11 min at 37 °C. 100 µl of reconstituted chromogenic substrate was added to the plate and gentle tapping was used to mix the solutions before returning the plate to 37 °C for 6 min. 50 µl of the stop reagent, 25% acetic acid, was added and the plate was mixed again by gentle tapping. We measured optical density at 405 nm. The average absorbance of blank wells containing endotoxin-free water was subtracted from the other measurements. We plotted the optical density at 405 nm as a function of endotoxin units (EU) per ml based on the standards, and the LPS concentration in the BEV sample was calculated from the resulting curve. The specification sheet for our purified LPS reported a conversion factor of 3,000,000 EU mg^-1^, which we used to express the measured concentration in terms of LPS mass per BEV.

### Pro-inflammatory stimulation

To test the direct effects of bacterial products on barrier integrity, cells were treated with LPS (Sigma-Aldrich, L3012-5MG) or BEVs diluted in hECSR medium + 2% FBS added to the µSiM top well or a multiwell plate. As a positive control for barrier disruption, cells were treated with an equal parts mixture of tumor necrosis factor alpha (TNF-α) (R&D Systems, 210TA), interleukin-1 beta (IL-1β) (R&D Systems, 201-LB), and interferon-gamma (IFN-γ) (R&D Systems, 285IF) referred to as cytomix. To examine the effects of macrophage–derived conditioned media on barrier integrity, THP-1 M0 macrophages were first treated with LPS or BEVs in serum-free RPMI 1640 + 1% anti-anti + 1% sodium pyruvate overnight. The collected conditioned media was twice centrifuged at 3,000×g for 15 min at 4 °C to pellet out cells and cell debris and then stored at -80°C until experimentation. For device treatment, the THP-1 conditioned media was diluted 1:2 with hECSR and added to the µSiM top well. Devices were incubated with the stimulants for 16–18 h prior to assessments of endothelial activation and barrier integrity.

### Fluorescent staining

For extracellular basement membrane staining, live cells were incubated with Collagen IV Monoclonal Antibody (1042), Alexa Fluor™ 647 (diluted 1:100, Invitrogen, 51-9871-82), Fibronectin Monoclonal Antibody (FN-3), Alexa Fluor™ 488 (diluted 1:200, Invitrogen, 53-9869-82), and Laminin Polyclonal Antibody (diluted 1:100, Invitrogen, PA1-16730) for 2 h. For ICAM-1 staining, live cells were incubated with purified anti-human CD54 Antibody (diluted 1:100, BioLegend, 353102) for 15 min. Cells were then washed with PBS and fixed with 4% formaldehyde for 15 min at room temperature. After three PBS washes, cells were blocked with 10% normal goat serum (Thermo Scientific, 50062Z) for 10–30 min at room temperature and then treated with secondary antibodies. Goat Anti-Rabbit IgG Alexa Fluor™ 568 (diluted 1:200, Invitrogen, A-11011) was used to label laminin and Goat Anti-Mouse IgG Alexa Fluor™ 488 (diluted 1:200, Invitrogen, A-11001) was used to label ICAM-1.

For claudin-5 and platelet–derived growth factor receptor β (PDGFRβ) labeling, cells were fixed in ice cold methanol for 20 s prior to washing three times with PBS. 5% normal goat serum + 0.4% Triton X-100 was used to block for 20 min at room temperature. The cells were stained with Mouse Anti-Human Claudin-5 IgG1 Alexa Fluor™ 488 (diluted 1:200, Invitrogen, 352588) and Rabbit Anti-Human PDGFRβ IgG (diluted 1:100, Cell Signaling Technology, 3169) for 1 h. After washing three times with PBS, the cells were treated with Goat Anti-Rabbit IgG, Alexa Fluor™ 568 (diluted 1:200, Invitrogen, A-11011) for 1 h to label PDGFRβ.

Nuclei were labeled with DAPI (Thermo Fisher, D1306) diluted 1:400 in deionized water for 3 min at room temperature or Hoechst 33342 diluted 1:10,000 in PBS for 1 h at room temperature prior to three final washes with PBS.

For live/dead staining, 0.7 µl calcein AM and 2.5 µl ethidium homodimer-1 (both from LIVE/DEAD Viability/Cytotoxicity Kit, Invitrogen, L3224) were mixed with 2 ml hECSR medium and 30 µl of this solution was added to live cells in a 96-well plate following a PBS wash. After 15 minutes of incubation in the dark at room temperature, the wells were flooded with 150 µl hECSR medium and imaged.

Standard epifluorescence images were captured on a Leica DMI6000B microscope equipped with a Rolera EM-C2™ EMCCD Camera (QImaging) using MetaMorph software (Molecular Devices, San Jose, California). 10x or 20x objective lenses were used for image acquisition. Confocal images were captured using an Andor Spinning Disk Confocal microscope stage (Abingdon, UK) attached to a Nikon TiE microscope (Nikon Corporation, Tokyo, Japan) equipped with a SONA sCMOS camera (Andor Technology, Belfast, UK). A 40x objective lens was used to acquire image stacks with a 0.2 μm step between images.

To analyze ICAM-1 expression in fluorescence images, the average fluorescence intensity in the ICAM-1 channel was measured and divided by the number of nuclei counted in corresponding DAPI images to yield the ICAM-1 intensity per cell. To determine the number of BEVs per BMEC, the number of nuclei was counted in each image and used to calculate the total number of cells per well. The number of BEVs added to each well was then divided by this value.

### Lucifer yellow permeability assay

Fresh hECSR medium was exchanged in the top well and lower channel of the µSiM. 150 µg ml^-1^ lucifer yellow (Invitrogen, L453) diluted in hECSR medium was added to the top well and the µSiM was incubated at 37 °C, 5% CO_2_. After 1 h, media was aspirated from the top well to stop diffusion. A pipette tip containing 50 µl hECSR medium was inserted into one of the open ports leading to the µSiM’s lower channel, and 50 µl of solution was sampled from the opposite port. The sample was added to a flat-bottomed black 96-well plate along with a reference ladder prepared by serially diluting the lucifer yellow solution. Samples were excited at 428 nm, and fluorescence intensity at 536 nm was measured on an Infinite 200 plate reader (TECAN, Männedorf, Switzerland). The concentration of lucifer yellow dye in the sample was determined by comparing to the linear region of the reference ladder. The system permeability *P*_*s*_ was calculated from equation 1:

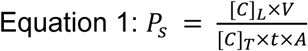

Where [*C*]_*L*_ is the dye concentration in the lower channel, *V* is the sample volume, [*C*]_*T*_ is the dye concentration added to the top well, *t* is the incubation time, and *A* is the surface area of the nanoporous membrane.

### Cytokine analysis

The concentrations of pro-inflammatory cytokines in conditioned media derived from THP-1 macrophages treated with BEVs or LPS were analyzed by Eve Technologies (Calgary, Alberta, Canada) using the Luminex® 200™ system (Luminex Corporation/DiaSorin, Saluggia, Italy) and Bio-Plex Manager™ software (Bio-Rad Laboratories Inc., Hercules, California, USA). Conditioned media samples were diluted 1:2 with PBS before shipping on dry ice. After arrival, the samples were run through the Human Cytokine Proinflammatory Focused 15-Plex Discovery Assay® Array (HDF15), which was designed to detect granulocyte-macrophage colony-stimulating factor (GM-CSF), IFN-γ, IL-1β, interleukin-1 receptor antagonist (IL-1RA), interleukin-2 (IL-2), interleukin-4, interleukin-5, interleukin-6 (IL-6), interleukin-8 (IL-8/CXCL8), interleukin-10, interleukin-12 p40, interleukin-12 p70, interleukin-13, monocyte chemoattractant protein-1 (MCP-1/CCL2), and TNF-α by capturing them with color-coded polystyrene beads conjugated to specific antibodies. A biotinylated detection antibody then bound to the captured cytokines as well as streptavidin-phycoerythrin. The beads were flowed through a dual-laser system that identified the type of cytokine based on the bead color and the amount of bound cytokine based on the phycoerythrin signal. To determine the concentration, the samples were compared to prepared standards of known concentrations. Our analysis excluded cytokines that fell below the assay’s detection range.

### Statistical analysis

Analysis was performed using GraphPad Prism software (GraphPad, La Jolla, California, USA). Data are presented as mean ± s.d. unless otherwise specified. For comparison between two conditions with a single independent variable, normally-distributed data, and no assumption of equal variance, Welch’s two-tailed t-test was used. For comparison between two conditions with a single independent variable, normally-distributed data, and equal variance, an unpaired two-tailed t-test was used. For comparisons between groups with a single independent variable and normally distributed data, one-way ANOVA with Dunnett’s test for multiple comparisons was used. *p* < 0.05 was considered significant.

## Results

### The µSiM supports a physiological BPB co-culture model

The BBB consists of several different cell types including brain microvascular endothelial cells and pericytes as well as a vascular basement membrane surrounding the blood vessels (Fig. 1a). To model the BBB, we used the µSiM platform. This 18 mm by 9 mm lab-on-chip device was previously developed by our group and validated for studies of the BBB (Fig. 1b).^18,19,49,51–53^ The central component of the µSiM is a porous silicon nitride membrane approximately 100 nm thick with abundant (10^10^ per cm^2^) ∼60 nm pores. This membrane was placed within an acrylic component above two layers of pressure sensitive adhesive and an underlying layer of cyclic olefin (Fig. 1c). The acrylic component formed an open top well above the membrane and provided ports to access a microfluidic channel located below the membrane. This enabled the introduction of cells and culture media to both sides of the porous membrane. The cyclic olefin layer provided a transparent window for high-quality imaging through the assembled µSiM device with inverted microscopes.

**Fig. 1:**
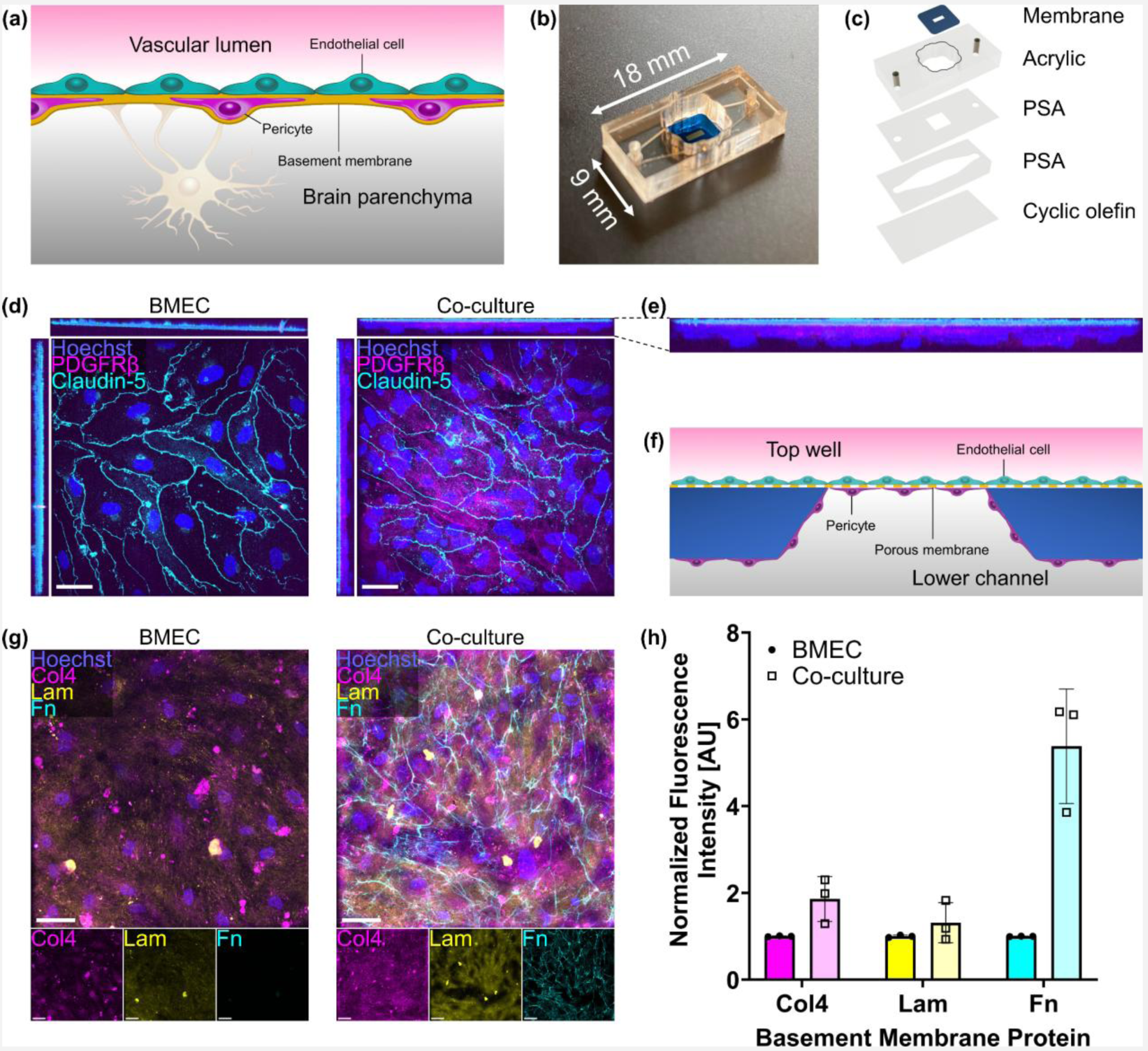
The μSiM supports a physiological BPB co-culture model. (**a**) Illustration of the BBB. (**b**) Dimensions of the µSiM. (**c**) µSiM assembly featuring an ultrathin silicon nitride porous membrane, acrylic top component, pressure sensitive adhesive (PSA) components, and bottom cyclic olefin layer. (**d**) Confocal images of a BMEC monoculture (left) and BMEC/BPLC co-culture (right) immunostained for nuclei (blue), PDGFRβ (magenta), and claudin-5 (cyan). X-axis projections are presented above each image, and y-axis projections to the left of each image. Scale bar = 50 µm. (**e**) Enlarged image of the x-axis projection of PDGFRβ and claudin-5 in a co-culture, demonstrating that BMECs were located above the BPLCs. (**f**) Illustration of cell seeding orientation, which matches the observation in panel (e). (**g**) Images of a BMEC monoculture (left) and BMEC/BPLC co-culture (right) immunostained for nuclei (blue), collagen IV (col4, magenta), laminin (lam, yellow), and fibronectin (fn, cyan). Images from the individual fluorescence channels are presented below the composite images. Scale bar = 50 µm. (**h**) Barplot of fluorescence intensities of collagen IV, laminin, and fibronectin normalized to the BMEC monoculture condition. Error bars = s.d.

To confirm that our co-culture model was consistent with previous µSiM work,^18^ we immunostained BPLCs on the underside of the membrane and BMECs on top of the membrane. BMEC monocultures demonstrated expression of claudin-5 tight junctions, while co-cultures possessed claudin-5 as well as PDGFRβ-positive BPLCs (Fig. 1d). Confocal microscopy yielded an x-axis projection which demonstrated that the claudin-5-positive BMECs were located above the PDGFRβ-positive BPLCs (Fig. 1e), which appropriately matched the seeding orientation that was selected to model the wall of brain blood vessels (Fig. 1f). This co-culture was termed the µSiM-BPB.

To examine whether the cells were able to manufacture their own basement membrane, we immunostained collagen IV, laminin, and fibronectin (Fig. 1g). Collagen IV and fibronectin were also utilized to coat the membranes prior to cell seeding. Cell-free coated membranes were immunostained, demonstrating punctate collagen IV that did not resemble the collagen IV deposited by cells, nor was there substantial labeling of fibronectin (See Fig. S1). Compared to BMEC monocultures, co-cultures had higher deposition of collagen IV and fibronectin (Fig. 1h). There was also a small increase in laminin deposition. It appeared that BPLCs contributed to basement membrane production, corroborating previous studies.^18,19^

### LPS and BEVs upregulate BMEC expression of ICAM-1

BEVs were isolated from *E. coli* strain CFT073 [WAM2267] by ultracentrifugation. Our BEV preparations were unable to form new bacterial colonies, confirming that all live *E. coli* were successfully removed. NTA was used to measure the concentration of recovered BEVs as well as their size distribution (Fig. 2a). We obtained a BEV concentration of (mean ± s.e.m.) 7.41×10^11^

**Fig. 2:**
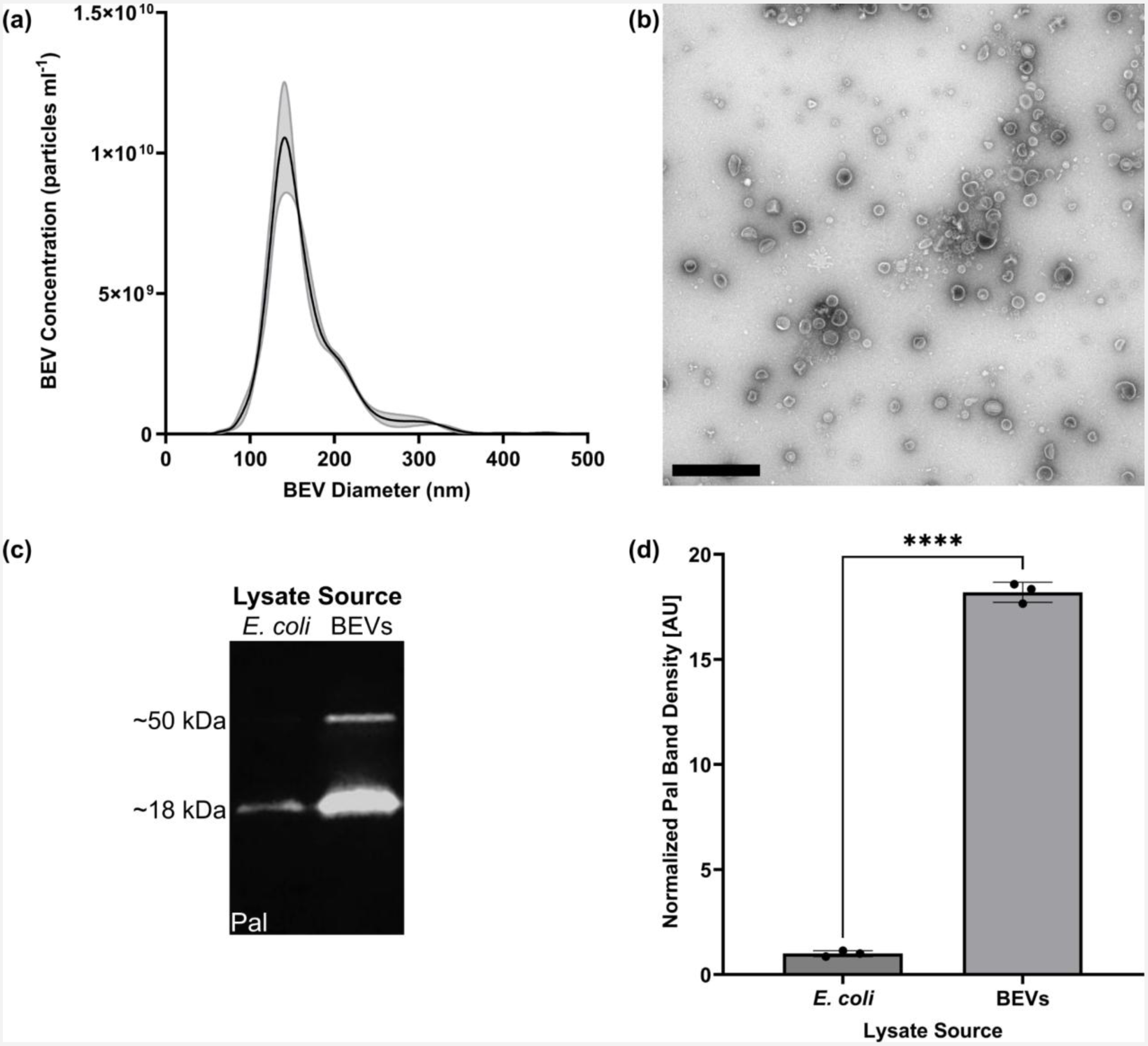
BEVs derived from *E. coli* strain CFT073 [WAM2267] demonstrate typical characteristics. (**a**) Representative NTA plot of BEV particle concentrations and size distribution. Error envelope = s.e.m. (**b**) TEM image of BEVs. Scale bar = 1 µm. (**c**) Fluorescent western blot of Pal protein from *E. coli* and BEV lysates. 1 µg total protein was added to each lane. This image was cropped from a single gel image; for the complete original western blot image, see Fig. S2. (**d**) Barplot of Pal band density at ∼18 kDa on western blot, normalized to the average band density measured in *E. coli* whole cell lysates. Error bars = s.d. Performed Welch’s two-tailed t-test. ✱✱✱✱ = *p* < 0.0001.

± 2.53×10^10^ particles ml^-1^ in our sample preparation, which corresponded to 9.88×10^9^ ± 3.37×10^8^ particles ml^-1^ in the *E. coli*-conditioned media prior to our isolation. The mean BEV diameters were 164.0 ± 1.1 nm. TEM demonstrated recovery of particles with morphologies typical of BEVs (Fig. 2b).^64^ Fluorescent western blot of equal amounts of total protein demonstrated that 18 kDa Pal was enriched in BEV lysates compared to *E. coli* whole cell lysates (Fig. 2c,d), which was expected since Pal is an outer membrane component that is typically packaged in *E. coli*–derived BEVs.^63,65^ An additional Pal band was observed at 50 kDa in BEV lysates but not *E. coli* lysates (Fig. 2c), which may correspond to complexes formed between Pal and outer membrane protein A.^66^ Previous work has demonstrated that *E. coli*–derived BEVs contain LPS that can initiate inflammation-associated processes.^32,33,35–37,67^ Endotoxin quantification demonstrated that our *E. coli*–derived BEVs contained 0.69 ± 0.01 fg LPS per particle.

To confirm that pure LPS and *E. coli*–derived BEVs could stimulate BMECs to upregulate ICAM-1, we seeded BMECs in multiwell plates and subjected them to treatment for 16–17 h. This treatment time was selected after determining that BMEC ICAM-1 expression increased over 10 h and remained elevated for up to 24 h after initial exposure to LPS (See Fig. S3). Immunostaining revealed increases in ICAM-1 on the BMEC surfaces in response to both LPS and BEV treatment (Fig. 3). 10 ng ml^-1^ LPS was able to significantly increase the ICAM-1 fluorescence signal, whereas 0.5 ng ml^-1^ LPS induced a non-significant increase (Fig. 3b). BEV treatment upregulated ICAM-1 expression at concentrations of both 1×10^6^ and 1×10^8^ particles ml^-1^ (Fig. 3d). This corresponded to (mean ± s.d.) 5.55 ± 0.73 BEVs per BMEC and 531.56 ± 47.15 BEVs per BMEC, respectively. Based on our measurements of the amount of LPS per particle, these treatments equated to 0.69 ng ml^-1^ and 69 ng ml^-1^ LPS, which were within the same order of magnitude as our LPS-only experiments. These experiments were conducted in media supplemented with 2% FBS to provide a source of LPS-binding protein (LBP) to aid BMEC sensing of LPS. In previous work, our lab and others have demonstrated that LPS mediates interactions between *E. coli*–derived BEVs and endothelial cells.^35–37,67^ Removal of FBS eliminated BMEC ICAM-1 upregulation in response to the BEVs (See Fig. S4).

**Fig. 3:**
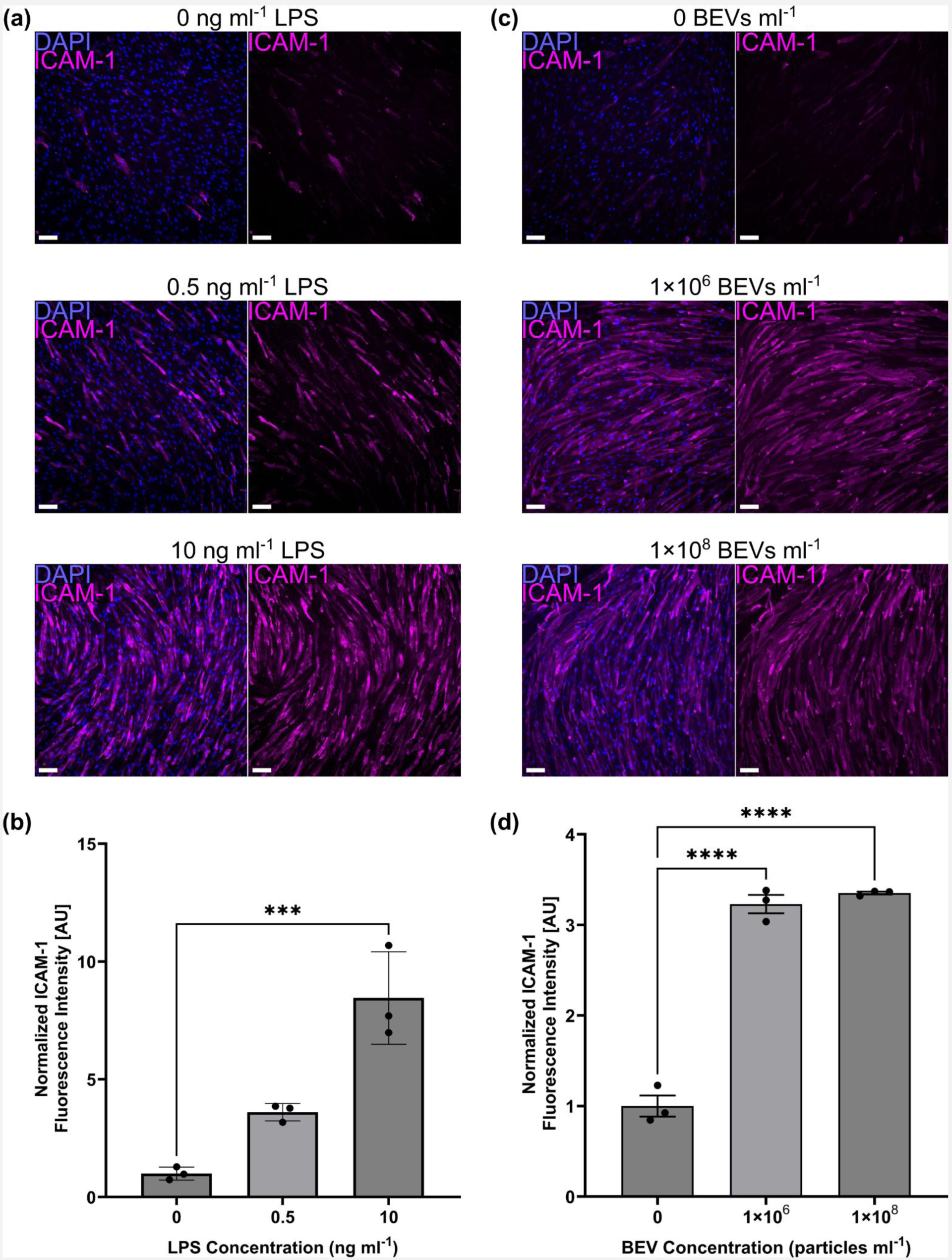
LPS and BEVs upregulate BMEC expression of ICAM-1. (**a**) Immunofluorescence images of BMECs in a multiwell plate stained for nuclei (blue) and ICAM-1 (magenta) following 16–17 h treatment with LPS. (**b**) Barplot of fluorescence intensity of ICAM-1 divided by cell count and normalized to the 0 ng ml^-1^ LPS condition. (**c**) Immunofluorescence images of BMECs in a multiwell plate stained for nuclei (blue) and ICAM-1 (magenta) following 16–17 h treatment with BEVs. (**d**) Barplot of fluorescence intensity of ICAM-1 divided by cell count and normalized to the 0 BEVs ml^-1^ condition. Scale bars = 100 µm. Error bars = s.d. Note that the y-axis scales of panels (b) and (d) are not comparable due to minor variations in staining and imaging parameters between the two experiments. Performed one-way ANOVA with Dunnett post-hoc test. ✱✱✱ = *p* < 0.001, ✱✱✱✱ = *p* < 0.0001.

### Neither LPS nor BEVs disrupt the BMEC barrier at the assayed treatment times and concentrations

To examine the effects of LPS and BEVs on barrier integrity, we established BMEC and BPLC co-cultures in µSiM devices and stained them for tight junction protein claudin-5 following 16–17 h treatment with each stimulant. At the concentrations and time points that we tested, there was no pronounced change in the fluorescence signal of claudin-5 (Fig. 4a,b). We also tested a pro-inflammatory “cytomix” consisting of equal parts TNF-α, IL-1β, and IFN-γ as a positive control for barrier disruption. This mixture mimics part of the cytokine storm that occurs during sepsis.^4^ At a concentration of 0.5 ng ml^-1^ (approximately 0.17 ng ml^-1^ of each individual cytokine), cytomix treatment caused substantial reduction of claudin-5 staining, indicating a loss of tight junction integrity (Fig. 4c).

**Fig. 4:**
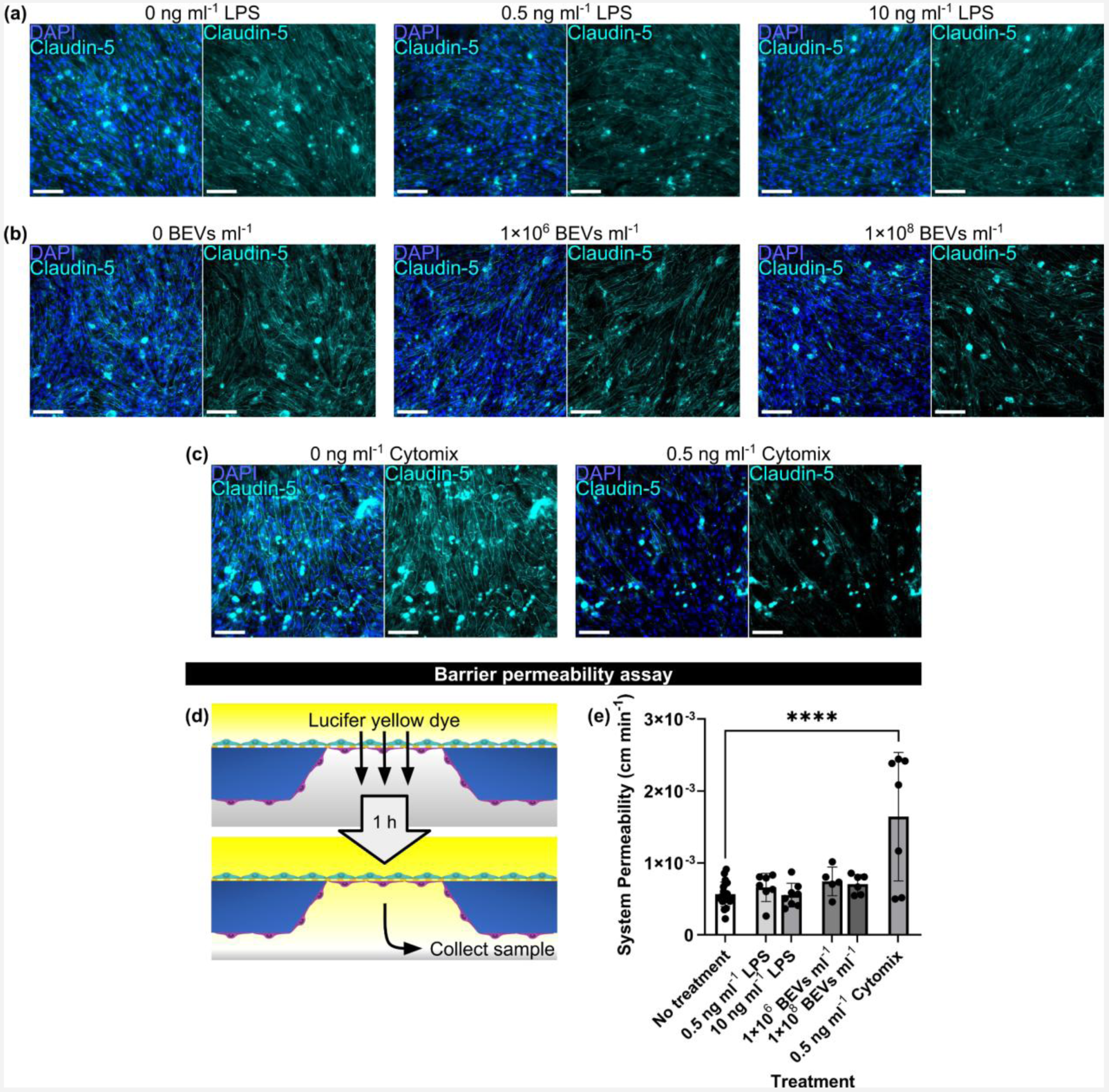
Neither LPS nor BEVs altered BMEC barrier properties. (**a**) Immunofluorescence images of BMEC/BPLC co-cultures immunostained for cell nuclei (blue) and claudin-5 (cyan) following 16–17 h treatment with LPS. (**b**) Immunofluorescence images of BMEC/BPLC co-cultures immunostained for cell nuclei (blue) and claudin-5 (cyan) following 16–17 h treatment with *E. coli*–derived BEVs. (**c**) Immunofluorescence images of BMEC/BPLC co-cultures immunostained for cell nuclei (blue) and claudin-5 (cyan) following 16–17 h treatment with a pro-inflammatory cytomix consisting of equal proportions of TNF-α, IL-1β, and IFN-γ. Cytomix mimics the effects of a cytokine storm and serves as a positive control for barrier disruption. (**d**) Illustration of permeability assay. Lucifer yellow dye (457 Da) was added to the µSiM top well. After 1 h, media was sampled from the lower channel so that its fluorescence intensity could be measured and compared to a prepared lucifer yellow standard curve with known concentrations. (**e**) Barplot of µSiM co-culture system permeability to lucifer yellow dye. Scale bars = 100 µm. Error bars = s.d. Performed one-way ANOVA with Dunnett post-hoc test. ✱✱✱✱ = *p* < 0.0001.

Permeability of the µSiM to lucifer yellow, a 457 Da dye molecule, was assessed by adding the dye to the µSiM top well and sampling from the lower channel after 1 h (Fig. 1d). Without treatment, the µSiM co-culture model had a system permeability of 5.64×10^-4^ ± 1.69×10^-4^ cm min^-1^. There was no significant increase in the average system permeability after LPS or BEV treatment compared to the untreated condition (6.59×10^-4^ ± 1.97×10^-4^ cm min^-1^ for 0.5 ng ml^-1^ LPS, 5.51×10^-4^ ± 1.66×10^-4^ cm min^-1^ for 10 ng ml^-1^ LPS, 7.43×10^-4^ ± 2.0×10^-4^ cm min^-1^ for 1×10^6^ BEVs ml^-1^, and 7.05×10^-4^ ± 1.34×10^-4^ cm min^-1^ for 1×10^8^ BEVs ml^-1^). Cytomix treatment resulted in a system permeability of 1.64×10^-3^ ± 8.93×10^-4^ cm min^-1^, which was the only condition that was significantly higher than the baseline permeability value (Fig. 4e).

### THP-1 macrophages are responsive to BEVs

To examine immune system interactions with BEVs, we used the THP-1 human monocyte cell line. Following 48 h treatment with 100 ng ml^-1^ phorbol 24-myristate 13-acetate, the THP-1s took on the phenotype of unpolarized macrophages. These cells were treated with 1×10^8^ BEVs ml^-1^ for 1, 2, 4, 8, 16, and 24 h. They showed a marked upregulation of ICAM-1 after 8 h, and this remained elevated through the 24 h timepoint (See Fig. S5). A dose-dependent relationship was also established after 16 h treatment, with increases in ICAM-1 signal starting at a concentration of 1×10^6^ BEVs ml^-1^ (See Fig. S6).

We intended to evaluate the response of the BMECs to conditioned media derived from THP-1 macrophages treated with BEVs, but first needed to establish whether the BMECs could tolerate mixtures of the THP-1 conditioned media for 16–18 h (See Fig. S7a). Live/dead staining demonstrated that BMEC monolayers tended to remain intact when the media mixture included up to 50% of conditioned media from untreated THP-1s mixed with hECSR. However, small gaps appeared when the THP-1 conditioned media concentration was increased to 75%, and with 100% conditioned media large gaps appeared in the BMEC monolayer. Despite these gaps, viability remained between 97.1–98.4% across the various conditions, though it is possible that some dead cells were washed away prior to imaging. Since the BMEC monolayer remained intact with a 50% mix of the THP-1 media, the baseline levels of ICAM-1 were examined under these conditions. No prominent difference in ICAM-1 upregulation was observed between the 50% mixture and 100% hECSR, though the 10 ng ml^-1^ LPS positive control did successfully cause increased ICAM-1 expression (See Fig. S7b).

### Conditioned medium from BEV-treated THP-1 macrophages disrupts the BMEC barrier

Finally, we modeled sepsis-like activation by treating BMECs with conditioned medium from cultures of THP-1 macrophages dosed for 24 h with 1×10^8^ BEVs ml^-1^. Following 16–17 h exposure to a 50% mixture of the conditioned medium, we immunostained BMEC monocultures for ICAM-1. ICAM-1 fluorescence intensity per cell was significantly increased when the conditioned medium was derived from THP-1s that were treated with BEVs compared to THP-1s that received no treatment (Fig. 5a,b). When the same conditioned medium was added to BMEC and BPLC co-cultures in µSiM devices for 16–17 h, we observed gaps in claudin-5 expression of the BMEC monolayer (Fig. 5c). These gaps appeared larger and more numerous in devices that received conditioned medium from THP-1s that were treated with BEVs. In addition, the lucifer yellow assay demonstrated that the co-cultures had significantly increased permeability when exposed to conditioned medium from BEV-treated THP-1s (9.59×10^-4^ ± 2.71×10^-4^ cm min^-1^) compared to untreated THP-1s (5.68×10^-4^ ± 2.01×10^-4^ cm min^-1^) (Fig. 5d).

**Fig. 5:**
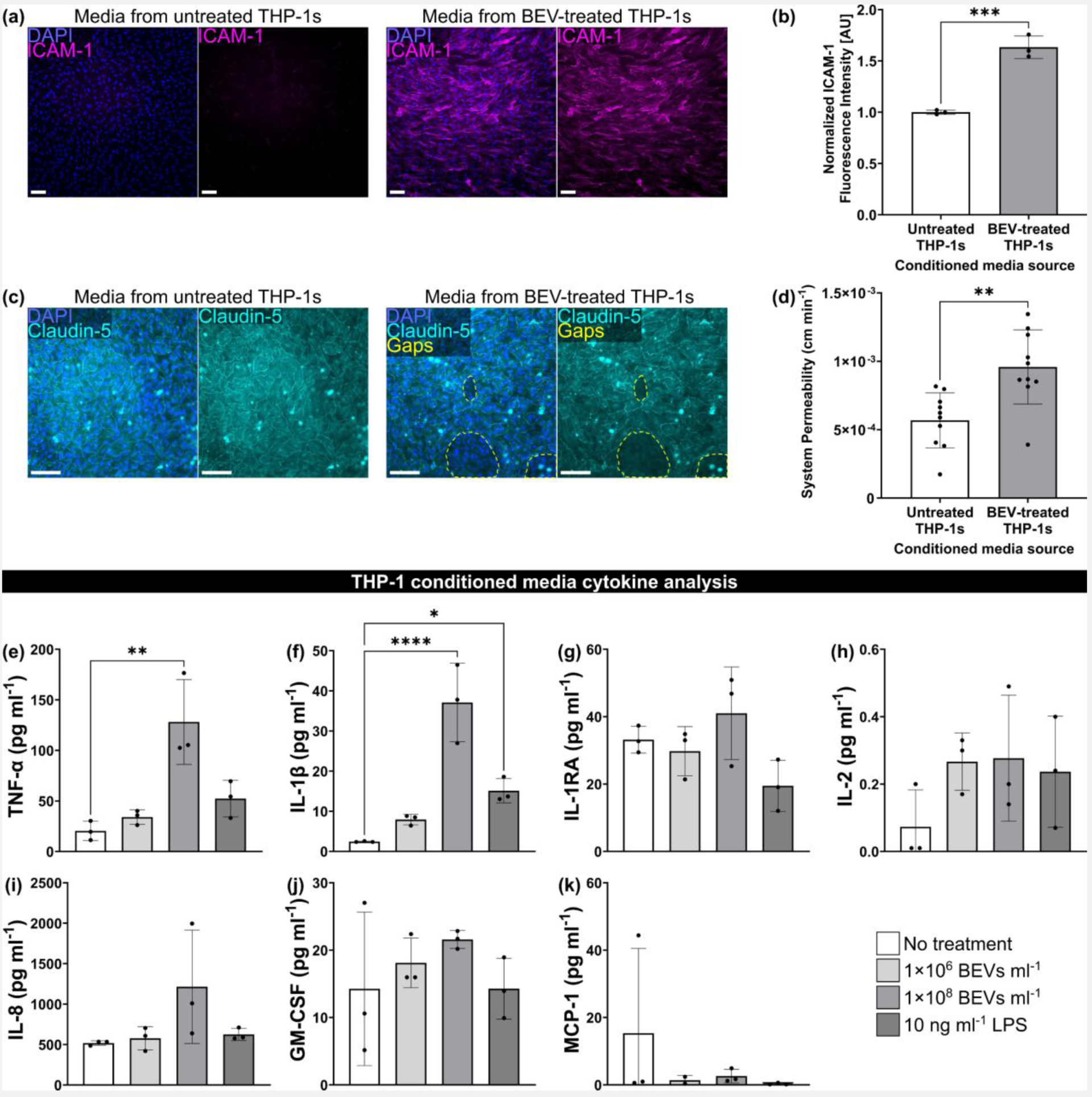
Conditioned medium from BEV-treated macrophages disrupts the barrier. (**a**) Immunofluorescence images of BMECs in a multiwell plate stained for nuclei (blue) and ICAM-1 (magenta) following 16–17 h treatment with 50% hECSR medium + 50% conditioned medium from THP-1 macrophages. The THP-1s had been treated with or without 1×10^8^ BEVs ml^-1^ for 24 h prior to collection of the conditioned medium. (**b**) Barplot of fluorescence intensity of ICAM-1 divided by cell count and normalized to the BMECs exposed to conditioned medium from untreated THP-1s. Performed unpaired two-tailed t-test. (**c**) Immunofluorescence images of BMEC/BPLC co-cultures immunostained for cell nuclei (blue) and claudin-5 (cyan) following 16–17 h treatment with 50% hECSR medium + 50% conditioned medium from THP-1 macrophages. Gaps in claudin-5 staining are circled with dotted lines (yellow). (**d**) Barplot of µSiM co-culture system permeability to lucifer yellow dye. Performed unpaired two-tailed t-test. (**e**)–(**k**) Cytokine analysis of conditioned medium from THP-1 macrophages that was collected following 24 h treatment with or without 1×10^6^ BEVs ml^-1^, 1×10^8^ BEVs ml^-1^, or 10 ng ml^-1^ LPS. Performed one-way ANOVA with Dunnett post-hoc test for each cytokine. (**e**) Barplot of TNF-α concentration. (**f**) Barplot of IL-1β concentration. (**g**) Barplot of IL-1RA concentration. (**h**) Barplot of IL-2 concentration. (**i**) Barplot of IL-8 concentration. (**j**) Barplot of GM-CSF concentration. (**k**) Barplot of MCP-1 concentration. Scale bars = 100 µm. Error bars = s.d. ✱ = *p* < 0.05, ✱✱ = *p* < 0.01, ✱✱✱ = *p* < 0.001, ✱✱✱✱ = *p* < 0.0001.

Concentrations of pro-inflammatory cytokines released by THP-1s into conditioned medium were quantified by Eve Technologies (Calgary, Alberta, Canada) (Fig. 5e–k). We examined two concentrations of BEVs, 1×10^6^ BEVs ml^-1^ and 1×10^8^ BEVs ml^-1^, as well as 10 ng ml^-1^ LPS. TNF-α demonstrated a significant increase from 20.47 ± 9.49 pg ml^-1^ to 128.2 ± 41.93 pg ml^-1^ after treatment with 1×10^8^ BEVs ml^-1^, and also trended upward after treatment with 1×10^6^ BEVs ml^-1^ and 10 ng ml^-1^ LPS, which yielded concentrations of 34.10 ± 7.15 pg ml^-1^ and 52.39 ± 18.21 pg ml^-1^, respectively (Fig. 5e). IL-1β, which had a concentration of 2.44 ± 0.12 pg ml^-1^ at baseline, demonstrated significant increases to 37.08 ± 9.77 pg ml^-1^ with 1×10^8^ BEVs ml^-1^ and 15.11 ± 3.04 pg ml^-1^ with 10 ng ml^-1^ LPS, as well as a non-significant increase to 7.92 ± 1.29 pg ml^-1^ with 1×10^6^ BEVs ml^-1^ treatment (Fig. 5f). While there were no significant differences between conditions for the other analyzed cytokines, the average concentration following treatment with 1×10^8^ BEVs ml^-1^ was consistently higher than the baseline (7.85 pg ml^-1^ higher for IL-1RA, 0.20 pg ml^-1^ higher for IL-2, 695.8 pg ml^-1^ higher for IL-8, and 7.31 pg ml^-1^ higher for GM-CSF). The only exception was MCP-1, though the higher average for the baseline was mainly driven by one datapoint. In addition to the cytokines displayed in Fig. 5, the analysis also included IL-6. IL-6 was generally too low to detect in our samples, but did appear following treatment with 1×10^8^ BEVs ml^-1^ (0.54 ± 0.36 pg ml^-1^ IL-6) and in a single replicate for the 10 ng ml^-1^ LPS condition (0.11 pg ml^-1^ IL-6). In summary, 24 h treatment of THP-1 macrophages with 1×10^8^ BEVs ml^-1^ enhanced the release of various pro-inflammatory cytokines. This increase could explain the changes in BMEC ICAM-1 expression and increased permeability to lucifer yellow dye.

## Discussion

BEVs are increasingly recognized as active contributors to both the initiation and persistence of sepsis. Treatment with antibiotics, which is currently the best defense against sepsis, can promote BEV release from bacteria, and it is possible that BEVs could linger in circulation following the clearance of the initial pathogenic infection.^63,65,68^ Various studies have demonstrated that BEVs promote a pro-inflammatory response in immune cells^38,39^ and may also destabilize the BBB.^42–44^ However, few, if any, studies have integrated pathogenic BEVs, the BBB, and immune interactions in an *in vitro* model mimicking human-specific physiology. This study sought to address this gap by examining the direct effects of BEVs on BBB integrity compared to indirect effects that were mediated by BEV–immune interactions.

Before conducting our BEV experiments, we focused on validation of our BBB model. The µSiM has previously been used to study BPB models, but we recognized the importance of confirming reproducibility, especially since this has been a challenge across the tissue chip community.^18,19^ To verify that our BMECs and BPLCs were situated in the correct seeding orientation, we immunostained claudin-5 and PDGFRβ. The latter is a receptor that is important for pericyte recruitment by endothelial cells,^69^ while the former is a key tight junction protein.^6^ Claudin-5 is primarily found in the brain and lungs, though it is also expressed by endothelial cells in the kidneys, liver, and skin.^70^ Its consistent appearance in our BMECs helped reinforce our decision to use these cells in our model, especially since other common model brain endothelial cells such as hCMEC/D3s reportedly demonstrate low claudin-5 expression and consequently have poor barrier integrity.^71^

The inclusion of BPLCs was previously observed to enhance barrier protection during pro-inflammatory challenge, indicating that a co-culture model was superior to BMECs alone when investigating inflammation.^18^ Furthermore, we observed that BPLCs were important for basement membrane deposition, as observed in prior studies.^18,19^ Collagen IV and laminin are considered the two largest components of the basement membrane,^15^ and while they appeared to be produced by our BMECs, it seemed that BPLCs also contributed to their production. The BPLCs also secreted large amounts of fibronectin, which may help contribute to tight junction expression by the endothelium.^72^ However, high fibronectin content in the basement membrane is typically associated with development or disease.^73^ It is possible that the quantity of fibronectin in our model would decrease given a longer maturation time.

A potential limitation of this study was the omission of astrocytes, which are another important component of the BBB. Previous µSiM studies have incorporated normal human astrocytes,^48,53^ but these cells have not yet been tested with BEVs. In murine models, BEVs derived from the gut bacteria of Alzheimer’s patients as well as BEVs secreted by *Helicobacter pylori* have caused astrocyte activation.^40,74^ It is probable that human astrocytes would react to treatment with our *E. coli*–derived BEVs, which would lead to greater neuroinflammation.

For this study, BEVs were isolated from *E. coli* by ultracentrifugation. This technique separates components from suspension based on mass. *E. coli* cells and larger debris were removed during low speed benchtop centrifugation and syringe filtration prior to the ultracentrifugation step, and the inability of our BEV preparations to form new bacterial colonies confirmed successful elimination of live *E. coli*. It is unlikely that smaller factors, such as 18–19 kDa free Pal protein, would pellet with the BEVs due to their low molecular weights.^75^ Ultracentrifugation is considered the “gold-standard” technique for extracellular vesicle isolation,^76^ and our NTA and TEM data show reasonable agreement with previously reported BEV characteristics.^34,64^ Other BEV isolation techniques may achieve higher analytical purity. However, excessive purification could come at the cost of physiological relevance in tissue chip systems, as it can strip away co-isolated bacterial components such as flagella, pili, and outer membrane fragments that are likely to be present in patient circulation following antibiotic treatment. Our laboratory is currently running a comparison between several isolation techniques including ultracentrifugation, ion-exchange chromatography, density gradient purification, and tangential flow filtration to better understand the differences in obtained BEV sample quality.

BEVs derived from pathogenic bacteria carry various pro-inflammatory stimuli including outer membrane protein A, Pal, and LPS.^32,65,77^ We have previously observed that LPS is the predominant stimulatory component in BEVs from *E. coli* strain CFT073 [WAM2267].^67^ Based on our endotoxin quantification results, the concentrations of BEV-associated LPS in our endothelial stimulation experiments were the same order of magnitude as the corresponding high and low concentrations of pure LPS that were used for comparison. The lower LPS dose that we used, 0.5 ng ml^-1^, was based on higher-end concentrations of LPS measured in serum from sepsis patients,^78^ whereas the 10 ng ml^-1^ LPS dose reflected amounts commonly used *in vitro*.^79–81^

Currently, it is challenging to isolate BEVs from patient blood due to their similarities to human extracellular vesicles.^82^ It would be interesting, however, if future analytical techniques could determine the ratio of LPS in sepsis patient blood incorporated in BEVs compared to free LPS. Mammalian extracellular vesicles have demonstrated a half-life of 30–360 min in blood,^83^ which is many times greater than free LPS’ 2–4 min half-life in mice.^84^ It is therefore likely that BEVs would also persist longer than free LPS and therefore have more opportunities to initiate host pro-inflammatory responses. Even if all measured LPS in sepsis patient blood was associated with BEVs, our results show that doses at this concentration would not cause vascular disruption without immune contributions.

Surprisingly, even though both BEVs and pure LPS were capable of upregulating BMEC ICAM-1 expression in our experiments, neither seemed to have a pronounced effect on claudin-5 expression or on permeability to lucifer yellow dye. This indicates that while ICAM-1 is a good reporter of endothelial reactivity, it does not necessarily signify barrier failure. Research by Pritchard *et al*. and Nonaka *et al*. has shown that BEVs derived from *Porphyromonas gingivalis* can reduce tight junction expression and increase permeability of brain endothelial cells *in vitro*.^42,43^ The observed differences could be due to the distinct source species for the BEVs. While the *E. coli* CFT073 strain used in the present study was selected for its known virulence and its association with urosepsis,^85^ it is conceivable that *P. gingivalis*–derived BEVs could be more stimulatory. Alternatively, it is possible that our hiPSC–derived BMECs are more resilient than the hCMEC/D3s and primary–derived human brain microvascular endothelial cells used in the other studies.^42,43^ hCMEC/D3s tend to have poor claudin-5 expression and barrier integrity, as mentioned previously, and our lab has had limited success establishing tight barriers with primary human brain endothelial cells in comparison to our hiPSC–derived BMECs (data not shown). Another possibility is that the inclusion of BPLCs made our model more resistant to BEV exposure, though Pritchard *et al*. included primary pericytes and astrocytes in a triculture with their brain endothelial cells.^42^ However, their cells were cultured across a traditional culture insert whose relatively high thickness may have impeded cell-cell communication compared to the ultrathin membrane in the µSiM, and thus potentially reduced protective effects bestowed by the mural cells.^50^

While BEVs and pure LPS treatments failed to disrupt barrier function in the assessed time period, our “cytomix” positive control treatment caused pronounced loss of barrier integrity. The TNF-α, IL-1β, and IFN-γ cytokines that made up our cytomix cocktail are highly elevated during the cytokine storms in sepsis.^4^ The cytomix concentrations used were 2–8 times higher than the mean concentrations of those seen in sepsis patients.^86^ This ensured a positive control that defined the upper end of the assay’s dynamic range. Not seeing a direct effect of physiologic LPS or BEV concentrations on barrier function, we then investigated if cytokine-rich conditioned media from BEV-treated THP-1 macrophages could trigger barrier disruption.

Conditioned media from BEV-treated THP-1 macrophages caused BMEC ICAM-1 upregulation, claudin-5 disruption, and increased permeability in the µSiM-BPB model. While we have attributed these results to THP-1-secreted cytokines in the conditioned media, it is possible that some residual BEVs were also present. However, this conditioned media was serum-free, and our experiments indicated that the BMECs required serum in order to respond to BEVs. We hypothesize that BMECs need serum protein LBP to bind LPS on the BEV surface for presentation to toll-like receptor 4.^87^ While THP-1 cells do not require serum proteins in order to detect BEVs, co-treatment with LBP would likely increase their sensitivity.^88,89^ Analysis of THP-1 monocyte and macrophage transcriptomes has revealed low, if any, LBP expression in these cells,^90^ so it is unlikely that they would be providing an alternate source of LBP for the endothelial cells. Thus, it is probable that the observed responses in the µSiM-BPB model were indeed caused by THP-1-secreted cytokines rather than residual BEVs in the conditioned media.

Regarding pro-inflammatory cytokines, we observed significant increases in TNF-α and IL-1β production by THP-1 macrophages in response to BEV treatment, as well as nonsignificant increases in IL-1RA, IL-2, IL-8, GM-CSF, and IL-6. Meganathan *et al*. similarly exposed THP-1 macrophages to BEVs isolated from poultry farm dust and saw increased TNF-α, IL-1β, IL-8, and IL-6.^38^ Other studies have confirmed that BEVs derived from various bacterial species, including *Pseudomonas aeruginosa*, *P. gingivalis*, *Treponema denticola*, *Tannerella forsythia*, and *E. coli*, are capable of causing THP-1s to increase production of pro-inflammatory cytokines.^91–93^ In the context of our model, this process recreated the disruptive cytokine storm that is observed during sepsis.

In the future, we plan to introduce monocytes directly into the µSiM-BPB model. It has long been understood that physiological fluid flow can influence monocyte arrest on endothelial cells as well as transmigration through the vascular wall.^94^ While the µSiM-BPB model was cultured under static conditions in the present study, a modular component has been developed allowing the addition of shear flow across the BMEC culture surface.^18,53,95^ By applying this technology in the context of BEV exposure, we may determine whether flow accentuates or diminishes endothelial and monocyte sensitivity to BEVs. We already observed the ability of BEVs to markedly upregulate BMEC ICAM-1 expression, so it is likely that monocyte adhesion to the endothelium would be increased following BEV pre-treatment of the µSiM-BPB. It would also be useful to examine the effects of pre-treating the monocytes with BEVs before adding them to the model. This would reveal whether BEV upregulation of endothelial ICAM-1 is necessary for monocyte recruitment or if signaling factors from the activated monocytes are sufficient to upregulate endothelial ICAM-1 to then provide adhesion sites for the monocytes. Finally, it is vital to investigate monocyte transmigration following BEV treatment. If exposure to pathogenic BEVs promotes monocyte infiltration into the brain, neuroinflammation will likely be worsened and sepsis sequelae will be exacerbated. After exploring these possibilities, we are interested in utilizing the µSiM-BPB model to explore therapeutic interventions that could dampen the immune response in the brain and reduce barrier disruption.

## Conclusions

The most important takeaway from the present study is that the examination of BEV interactions with the BBB should not be performed in oversimplified systems lacking an immune component. While this is hardly the first study to examine how BEVs modulate immune cell secretions, to the best of our knowledge, it is the first to apply these secretions to an *in vitro* BBB model to examine downstream effects. Notably, barrier disruption due to treatment with conditioned media from BEV-treated macrophages was more pronounced than direct BBB exposure to bacterial products, supporting the notion that the most significant barrier damage during sepsis is due to host-secreted cytokines. The inclusion of leukocyte–derived components should strengthen the predictive power of BBB models and in the future will hopefully aid development of therapeutic interventions to mitigate BEV-associated BBB disruption and improve sepsis outcomes.

## Author contributions

LPW, JLM, and TRG Conceptualization; LPW Formal Analysis; LVM, JLM, and TRG Funding Acquisition; LPW Investigation; LPW, PT, MAT, MCM, LVM, JLM, and TRG Methodology; LPW, JLM, and TRG Project Administration; MAT, MCM, LVM, JLM, and TRG Resources; JLM and TRG Supervision; LPW, PT, MAT, MCM Validation; LPW Visualization, LPW Writing – Original Draft Preparation.

## Data availability

The majority of relevant data for this study are presented in the manuscript and supporting information. Fluorescence data, NTA data, western blot data, LPS quantification data, permeability data, and cytokine concentration data are available on Figshare (DOI: 10.6084/m9.figshare.31118659).

## Acknowledgements

Transmission electron microscopy was performed by the staff of the Electron Microscopy Resource, part of the Center for Advanced Research Technology at the University of Rochester Medical Center. Cytokine analysis was performed by Eve Technologies. Thank you to the High Content Image Core (University of Rochester) for assistance with fluorescent imaging and use of the Dragonfly Spinning Disk Confocal microscope. This work was supported by the National Institute of Allergy and Infectious Diseases (NIAID) of the National Institutes of Health under Award No. R21AI163782 to LVM and TRG, the National Heart, Lung, and Blood Institute (NHLBI) through grant R33HL154249 to JLM and TRG, the National Institute of General Medical Sciences (NIGMS) through grant R35GM153461 to TRG, and the National Institute of Aging (NIA) through grant U2CAG088071 to JLM and TRG. Additional support was provided by the Hank and Lynn Hopeman Foundation, also awarded to TRG. These funding sources did not influence the study design; collection, analysis, and interpretation of data; writing of the paper; or decision to submit for publication.

## References

1. Singer M, Deutschman CS, Seymour CW, et al. The Third International Consensus Definitions for Sepsis and Septic Shock (Sepsis-3). JAMA 2016; 315: 801–810.

2. Rhee C, Dantes R, Epstein L, et al. Incidence and Trends of Sepsis in US Hospitals Using Clinical vs Claims Data, 2009-2014. JAMA 2017; 318: 1241–1249.

3. Evans T. Diagnosis and management of sepsis. Clin Med 2018; 18: 146–149.

4. Fajgenbaum DC, June CH. Cytokine Storm. N Engl J Med 2020; 383: 2255–2273.

5. Mostel Z, Perl A, Marck M, et al. Post-sepsis syndrome – an evolving entity that afflicts survivors of sepsis. Mol Med 2019; 26: 6.

6. Cong X, Kong W. Endothelial tight junctions and their regulatory signaling pathways in vascular homeostasis and disease. Cell Signal 2020; 66: 109485.

7. Gabathuler R. Approaches to transport therapeutic drugs across the blood–brain barrier to treat brain diseases. Neurobiol Dis 2010; 37: 48–57.

8. Hall CN, Reynell C, Gesslein B, et al. Capillary pericytes regulate cerebral blood flow in health and disease. Nature 2014; 508: 55–60.

9. Armulik A, Genové G, Mäe M, et al. Pericytes regulate the blood–brain barrier. Nature 2010; 468: 557–561.

10. Bhowmick S, D’Mello V, Caruso D, et al. Impairment of pericyte-endothelium crosstalk leads to blood-brain barrier dysfunction following traumatic brain injury. Exp Neurol 2019; 317: 260–270.

11. Herland A, van der Meer AD, FitzGerald EA, et al. Distinct Contributions of Astrocytes and Pericytes to Neuroinflammation Identified in a 3D Human Blood-Brain Barrier on a Chip. PLOS ONE 2016; 11: e0150360.

12. Ayloo S, Lazo CG, Sun S, et al. Pericyte-to-endothelial cell signaling via vitronectin-integrin regulates blood-CNS barrier. Neuron. Epub ahead of print 15 March 2022. DOI: 10.1016/j.neuron.2022.02.017.

13. Thomsen MS, Birkelund S, Burkhart A, et al. Synthesis and deposition of basement membrane proteins by primary brain capillary endothelial cells in a murine model of the blood–brain barrier. J Neurochem 2017; 140: 741–754.

14. Caporarello N, D’Angeli F, Cambria MT, et al. Pericytes in Microvessels: From “Mural” Function to Brain and Retina Regeneration. Int J Mol Sci 2019; 20: 6351.

15. Leclech C, Natale CF, Barakat AI. The basement membrane as a structured surface – role in vascular health and disease. J Cell Sci 2020; 133: jcs239889.

16. Stratman AN, Malotte KM, Mahan RD, et al. Pericyte recruitment during vasculogenic tube assembly stimulates endothelial basement membrane matrix formation. Blood 2009; 114: 5091–5101.

17. Yamazaki Y, Shinohara M, Yamazaki A, et al. ApoE (Apolipoprotein E) in Brain Pericytes Regulates Endothelial Function in an Isoform-Dependent Manner by Modulating Basement Membrane Components. Arterioscler Thromb Vasc Biol 2020; 40: 128–144.

18. McCloskey MC, Ahmad SD, Widom LP, et al. Pericytes Enrich the Basement Membrane and Reduce Neutrophil Transmigration in an In Vitro Model of Peripheral Inflammation at the Blood–Brain Barrier. Biomater Res 2024; 28: 0081.

19. Trempel MA, Du Y, Widom LP, et al. Pericytes Repair Engineered Defects in the Basement Membrane to Restore Barrier Integrity in an in vitro Model of the Blood-Brain Barrier. Mater Today Bio 2025; 102361.

20. Osada T, Gu Y-H, Kanazawa M, et al. Interendothelial claudin-5 expression depends on cerebral endothelial cell-matrix adhesion by β(1)-integrins. J Cereb Blood Flow Metab Off J Int Soc Cereb Blood Flow Metab 2011; 31: 1972–1985.

21. Barichello T, Generoso JS, Collodel A, et al. The blood-brain barrier dysfunction in sepsis. Tissue Barriers 2021; 9: 1840912.

22. Erikson K, Tuominen H, Vakkala M, et al. Brain tight junction protein expression in sepsis in an autopsy series. Crit Care 2020; 24: 385.

23. Lawson C, Wolf S. ICAM-1 signaling in endothelial cells. Pharmacol Rep 2009; 61: 22–32.

24. Van De Stolpe A, Van Der Saag PT. Intercellular adhesion molecule-1. J Mol Med 1996; 74: 13–33.

25. Zhong H, Lin H, Pang Q, et al. Macrophage ICAM-1 functions as a regulator of phagocytosis in LPS induced endotoxemia. Inflamm Res 2021; 70: 193–203.

26. Trzeciak A, Lerman YV, Kim T-H, et al. Long-Term Microgliosis Driven by Acute Systemic Inflammation. J Immunol 2019; 203: 2979–2989.

27. Andonegui G, Zelinski EL, Schubert CL, et al. Targeting inflammatory monocytes in sepsis-associated encephalopathy and long-term cognitive impairment. JCI Insight 2018; 3: e99364.

28. Rudziak P, Ellis CG, Kowalewska PM. Role and Molecular Mechanisms of Pericytes in Regulation of Leukocyte Diapedesis in Inflamed Tissues. Mediators Inflamm 2019; 2019: 1–9.

29. Zeng H, He X, Tuo Q, et al. LPS causes pericyte loss and microvascular dysfunction via disruption of Sirt3/angiopoietins/Tie-2 and HIF-2α/Notch3 pathways. Sci Rep 2016; 6: 20931.

30. Effah CY, Ding X, Drokow EK, et al. Bacteria-derived extracellular vesicles: endogenous roles, therapeutic potentials and their biomimetics for the treatment and prevention of sepsis. Front Immunol 2024; 15: 1296061.

31. Brown L, Wolf JM, Prados-Rosales R, et al. Through the wall: extracellular vesicles in Gram-positive bacteria, mycobacteria and fungi. Nat Rev Microbiol 2015; 13: 620–630.

32. Beveridge TJ. Structures of Gram-Negative Cell Walls and Their Derived Membrane Vesicles. J Bacteriol 1999; 181: 4725–4733.

33. Hosseini-Giv N, Basas A, Hicks C, et al. Bacterial extracellular vesicles and their novel therapeutic applications in health and cancer. Front Cell Infect Microbiol 2022; 12: 962216.

34. Toyofuku M, Schild S, Kaparakis-Liaskos M, et al. Composition and functions of bacterial membrane vesicles. Nat Rev Microbiol 2023; 21: 415–430.

35. Laakmann K, Eckersberg JM, Hapke M, et al. Bacterial extracellular vesicles repress the vascular protective factor RNase1 in human lung endothelial cells. Cell Commun Signal 2023; 21: 111.

36. Soult MC, Lonergan NE, Shah B, et al. Outer membrane vesicles from pathogenic bacteria initiate an inflammatory response in human endothelial cells. J Surg Res 2013; 184: 458–466.

37. Kim JH, Yoon YJ, Lee J, et al. Outer Membrane Vesicles Derived from Escherichia coli Up-Regulate Expression of Endothelial Cell Adhesion Molecules In Vitro and In Vivo. PLoS ONE 2013; 8: e59276.

38. Meganathan V, Moyana R, Natarajan K, et al. Bacterial extracellular vesicles isolated from organic dust induce neutrophilic inflammation in the lung. Am J Physiol-Lung Cell Mol Physiol 2020; 319: L893–L907.

39. Wei S, Jiao D, Xing W. A rapid method for isolation of bacterial extracellular vesicles from culture media using epsilon-poly-L–lysine that enables immunological function research. Front Immunol 2022; 13: 930510.

40. Wei S, Peng W, Mai Y, et al. Outer membrane vesicles enhance tau phosphorylation and contribute to cognitive impairment. J Cell Physiol 2020; 235: 4843–4855.

41. Gong T, Chen Q, Mao H, et al. Outer membrane vesicles of Porphyromonas gingivalis trigger NLRP3 inflammasome and induce neuroinflammation, tau phosphorylation, and memory dysfunction in mice. Front Cell Infect Microbiol 2022; 12: 925435.

42. Pritchard AB, Fabian Z, Lawrence CL, et al. An Investigation into the Effects of Outer Membrane Vesicles and Lipopolysaccharide of Porphyromonas gingivalis on Blood-Brain Barrier Integrity, Permeability, and Disruption of Scaffolding Proteins in a Human in vitro Model. J Alzheimers Dis 2022; 86: 343–364.

43. Nonaka S, Kadowaki T, Nakanishi H. Secreted gingipains from Porphyromonas gingivalis increase permeability in human cerebral microvascular endothelial cells through intracellular degradation of tight junction proteins. Neurochem Int 2022; 154: 105282.

44. Kaisanlahti A, Salmi S, Kumpula S, et al. Bacterial extracellular vesicles – brain invaders? A systematic review. Front Mol Neurosci 2023; 16: 1227655.

45. Kangwantas K, Pinteaux E, Penny J. The extracellular matrix protein laminin-10 promotes blood–brain barrier repair after hypoxia and inflammation in vitro. J Neuroinflammation 2016; 13: 25.

46. Ding X, Sun X, Shen X, et al. Propofol attenuates TNF-α-induced MMP-9 expression in human cerebral microvascular endothelial cells by inhibiting Ca2+/CAMK II/ERK/NF-κB signaling pathway. Acta Pharmacol Sin 2019; 40: 1303–1313.

47. Khire TS, Salminen AT, Swamy H, et al. Microvascular Mimetics for the Study of Leukocyte–Endothelial Interactions. Cell Mol Bioeng 2020; 13: 125–139.

48. McCloskey MC, Kasap P, Ahmad SD, et al. The Modular µSiM: A Mass Produced, Rapidly Assembled, and Reconfigurable Platform for the Study of Barrier Tissue Models In Vitro. Adv Healthc Mater 2022; 11: 2200804.

49. McCloskey MC, Kasap P, Trempel M, et al. Use of the MicroSiM (µSiM) Barrier Tissue Platform for Modeling the Blood-Brain Barrier. J Vis Exp 2024; 65258.

50. Gholizadeh S, Carter RN, Allahyari Z, et al. Deciphering the role of physical proximity on endothelial-glial interactions using thickness gradient nanomembranes. Biophys J 2022; 121: 424a.

51. Mossu A, Rosito M, Khire T, et al. A silicon nanomembrane platform for the visualization of immune cell trafficking across the human blood–brain barrier under flow. J Cereb Blood Flow Metab 2019; 39: 395–410.

52. Hudecz D, McCloskey MC, Vergo S, et al. Modelling a Human Blood-Brain Barrier Co-Culture Using an Ultrathin Silicon Nitride Membrane-Based Microfluidic Device. Int J Mol Sci 2023; 24: 5624.

53. Chen K, Linares IM, Trempel MA, et al. Shear Conditioning Promotes Microvascular Endothelial Barrier Resilience in a Human BBB-on-a-Chip Model of Systemic Inflammation Leading to Astrogliosis. Adv Sci 2025; 12: e08271.

54. Nishihara H, Gastfriend BD, Kasap P, et al. Differentiation of human pluripotent stem cells to brain microvascular endothelial cell-like cells suitable to study immune cell interactions. STAR Protoc 2021; 2: 100563–100563.

55. Gastfriend BD, Stebbins MJ, Du F, et al. Differentiation of Brain Pericyte-Like Cells from Human Pluripotent Stem Cell-Derived Neural Crest. Curr Protoc 2021; 1: e21.

56. Engelhardt B, Vajkoczy P, Weller RO. The movers and shapers in immune privilege of the CNS. Nat Immunol 2017; 18: 123–131.

57. Kucharz K, Kristensen K, Johnsen KB, et al. Post-capillary venules are the key locus for transcytosis-mediated brain delivery of therapeutic nanoparticles. Nat Commun 2021; 12: 4121.

58. Su S-H, Song Y, Stephens A, et al. A tissue chip with integrated digital immunosensors: In situ brain endothelial barrier cytokine secretion monitoring. Biosens Bioelectron 2023; 224: 115030.

59. Seok J, Warren HS, Cuenca AG, et al. Genomic responses in mouse models poorly mimic human inflammatory diseases. Proc Natl Acad Sci 2013; 110: 3507–3512.

60. Taylor G. Animal models of respiratory syncytial virus infection. Vaccine 2017; 35: 469–480.

61. Emini Veseli B, Perrotta P, De Meyer GRA, et al. Animal models of atherosclerosis. Eur J Pharmacol 2017; 816: 3–13.

62. Burma NE, Leduc-Pessah H, Fan CY, et al. Animal models of chronic pain: Advances and challenges for clinical translation. J Neurosci Res 2017; 95: 1242–1256.

63. Torabian P, Singh N, Crawford J, et al. Effect of clinically relevant antibiotics on bacterial extracellular vesicle release from Escherichia coli. Int J Antimicrob Agents 2025; 65: 107384.

64. Wen M, Wang J, Ou Z, et al. Bacterial extracellular vesicles: A position paper by the microbial vesicles task force of the Chinese society for extracellular vesicles. Interdiscip Med 2023; 1: e20230017.

65. Michel LV, Gallardo L, Konovalova A, et al. Ampicillin triggers the release of Pal in toxic vesicles from Escherichia coli. Int J Antimicrob Agents 2020; 56: 106163.

66. Cascales E, Bernadac A, Gavioli M, et al. Pal Lipoprotein of *Escherichia coli* Plays a Major Role in Outer Membrane Integrity. J Bacteriol 2002; 184: 754–759.

67. Widom LP, Torabian P, Wojehowski AC, et al. Antibiotic treatment modulates *Escherichia coli*-derived bacterial extracellular vesicle (BEV) production and their capacity to upregulate ICAM-1 in human endothelial cells. Epub ahead of print 13 May 2025. DOI: 10.1101/2025.05.12.653517.

68. Ye C, Li W, Yang Y, et al. Inappropriate use of antibiotics exacerbates inflammation through OMV-induced pyroptosis in MDR Klebsiella pneumoniae infection. Cell Rep 2021; 36: 109750.

69. Darland DC, D’Amore PA. Blood vessel maturation: vascular development comes of age. J Clin Invest 1999; 103: 157–158.

70. Greene C, Hanley N, Campbell M. Claudin-5: gatekeeper of neurological function. Fluids Barriers CNS 2019; 16: 3.

71. Gericke B, Römermann K, Noack A, et al. A face-to-face comparison of claudin-5 transduced human brain endothelial (hCMEC/D3) cells with porcine brain endothelial cells as blood–brain barrier models for drug transport studies. Fluids Barriers CNS 2020; 17: 53.

72. Tilling T, Korte D, Hoheisel D, et al. Basement Membrane Proteins Influence Brain Capillary Endothelial Barrier Function In Vitro. J Neurochem 1998; 71: 1151–1157.

73. Thomsen MS, Routhe LJ, Moos T. The vascular basement membrane in the healthy and pathological brain. J Cereb Blood Flow Metab Off J Int Soc Cereb Blood Flow Metab 2017; 37: 3300–3317.

74. Park A-M, Tsunoda I. Helicobacter pylori infection in the stomach induces neuroinflammation: the potential roles of bacterial outer membrane vesicles in an animal model of Alzheimer’s disease. Inflamm Regen 2022; 42: 39.

75. Hellman J, Roberts JD, Tehan MM, et al. Bacterial Peptidoglycan-associated Lipoprotein Is Released into the Bloodstream in Gram-negative Sepsis and Causes Inflammation and Death in Mice. J Biol Chem 2002; 277: 14274–14280.

76. Saint-Pol J, Culot M. Minimum information for studies of extracellular vesicles (MISEV) as toolbox for rigorous, reproducible and homogeneous studies on extracellular vesicles. Toxicol In Vitro 2025; 106: 106049.

77. Skerniškytė J, Karazijaitė E, Lučiūnaitė A, et al. OmpA Protein-Deficient Acinetobacter baumannii Outer Membrane Vesicles Trigger Reduced Inflammatory Response. Pathogens 2021; 10: 407.

78. Marshall JC, Walker PM, Foster DM, et al. Measurement of endotoxin activity in critically ill patients using whole blood neutrophil dependent chemiluminescence. Crit Care 2002; 6: 342.

79. Henry CJ, Huang Y, Wynne A, et al. Minocycline attenuates lipopolysaccharide (LPS)-induced neuroinflammation, sickness behavior, and anhedonia. J Neuroinflammation 2008; 5: 15.

80. Puffenbarger RA, Boothe AC, Cabral GA. Cannabinoids inhibit LPS-inducible cytokine mRNA expression in rat microglial cells. Glia 2000; 29: 58–69.

81. De Vries HE, Moor ACE, Blom-Roosemalen MCM, et al. Lymphocyte adhesion to brain capillary endothelial cells in vitro. J Neuroimmunol 1994; 52: 1–8.

82. Arnold P, Boros FA, Mattner J, et al. Two Sides of the Same Coin—Mechanistic Insight, Diagnostic Application and Therapeutic Translation of Bacterial and Host-Derived Extracellular Vesicles. J Extracell Biol 2025; 4: e70093.

83. Karaman I, Pathak A, Bayik D, et al. Harnessing Bacterial Extracellular Vesicle Immune Effects for Cancer Therapy. Pathog Immun 2024; 9: 56–90.

84. Yao Z, Mates JM, Cheplowitz AM, et al. Blood-Borne Lipopolysaccharide Is Rapidly Eliminated by Liver Sinusoidal Endothelial Cells via High-Density Lipoprotein. J Immunol 2016; 197: 2390–2399.

85. Anfora AT, Halladin DK, Haugen BJ, et al. Uropathogenic *Escherichia coli* CFT073 Is Adapted to Acetatogenic Growth but Does Not Require Acetate during Murine Urinary Tract Infection. Infect Immun 2008; 76: 5760–5767.

86. Gharamti AA, Samara O, Monzon A, et al. Proinflammatory cytokines levels in sepsis and healthy volunteers, and tumor necrosis factor-alpha associated sepsis mortality: A systematic review and meta-analysis. Cytokine 2022; 158: 156006.

87. Dauphinee SM, Karsan A. Lipopolysaccharide signaling in endothelial cells. Lab Invest 2006; 86: 9–22.

88. Tobias PS, Soldau K, Kline L, et al. Cross-linking of lipopolysaccharide (LPS) to CD14 on THP-1 cells mediated by LPS-binding protein. J Immunol 1993; 150: 3011–3021.

89. Su GL, Simmons RL, Wang SC. Lipopolysaccharide Binding Protein Participation in Cellular Activation by LPS. Crit Rev Immunol 1995; 15: 201–214.

90. Liu T, Huang T, Li J, et al. Optimization of differentiation and transcriptomic profile of THP-1 cells into macrophage by PMA. PLOS ONE 2023; 18: e0286056.

91. Lee J, Yoon YJ, Kim JH, et al. Outer Membrane Vesicles Derived From Escherichia coli Regulate Neutrophil Migration by Induction of Endothelial IL-8. Front Microbiol 2018; 9: 2268.

92. Ge J, Liu Y, Wu T, et al. Outer membrane vesicles from Pseudomonas aeruginosa induce autophagy-regulated pyroptosis in THP-1 cells. Arch Microbiol 2025; 207: 54.

93. Cecil JD, O’Brien-Simpson NM, Lenzo JC, et al. Outer Membrane Vesicles Prime and Activate Macrophage Inflammasomes and Cytokine Secretion In Vitro and In Vivo. Front Immunol 2017; 8: 1017.

94. Weber KSC, Von Hundelshausen P, Clark-Lewis I, et al. Differential immobilization and hierarchical involvement of chemokines in monocyte arrest and transmigration on inflamed endothelium in shear flow. Eur J Immunol 1999; 29: 700–712.

95. Mansouri M, Ahmed A, Ahmad SD, et al. The Modular µSiM Reconfigured: Integration of Microfluidic Capabilities to Study In Vitro Barrier Tissue Models under Flow. Adv Healthc Mater 2022; 11: 2200802.

