## Supporting Information for "Bacterial extracellular vesicles indirectly destabilize a human stem cell–derived blood–brain barrier on-chip through pro-inflammatory stimulation of immune cells"

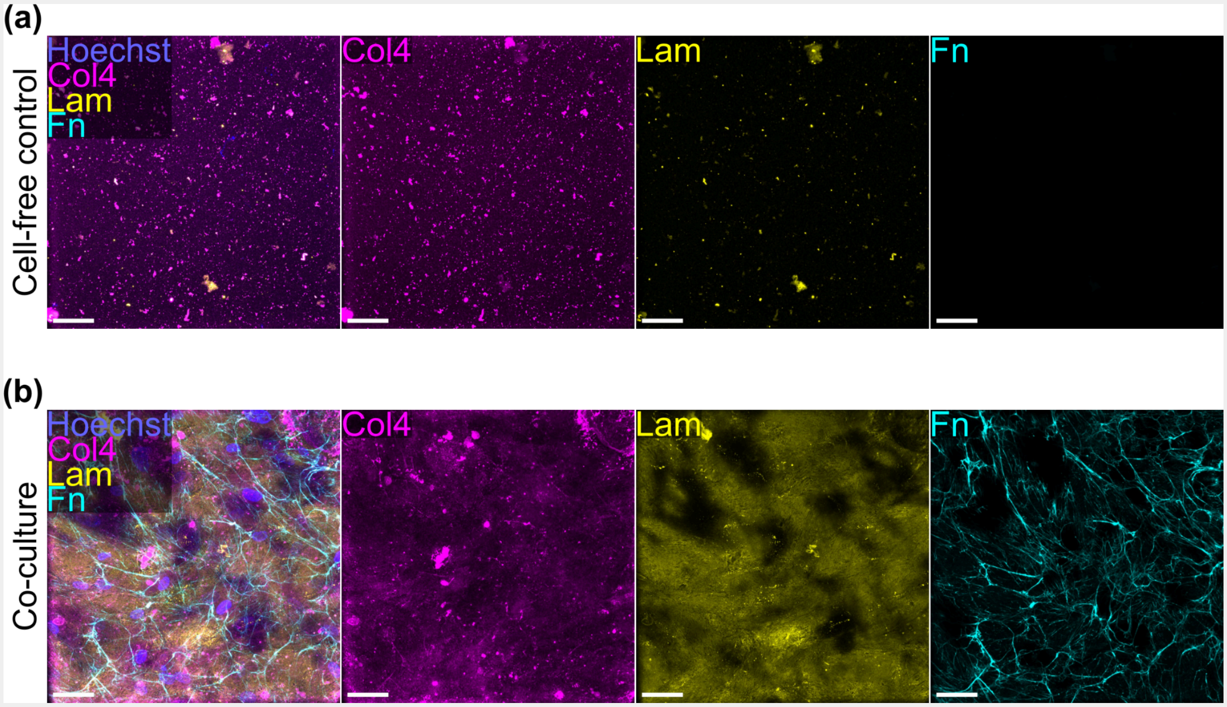


**Fig. S1: Coated cell-free membranes do not demonstrate substantial basement membrane staining compared to cell-seeded membranes.** (**a**) Images of a cell-free control membrane coated with collagen IV and fibronectin immunostained for cell nuclei (blue), collagen IV (col4, magenta), laminin (lam, yellow), and fibronectin (fn, cyan). Images from the individual fluorescence channels are presented to the right of the composite image. Collagen IV and laminin demonstrated punctate staining, and very little fibronectin was visible. (**b**) Images of a BMEC/BPLC co-culture labeled with the same stains as panel (a). Images from the individual fluorescence channels are presented to the right of the composite image. Collagen IV and laminin were less punctate and the appearance of organized structures was indicative that the proteins were deposited and organized by the cells. There was also substantial labeling of fibronectin networks. Scale bars = 50 μm.

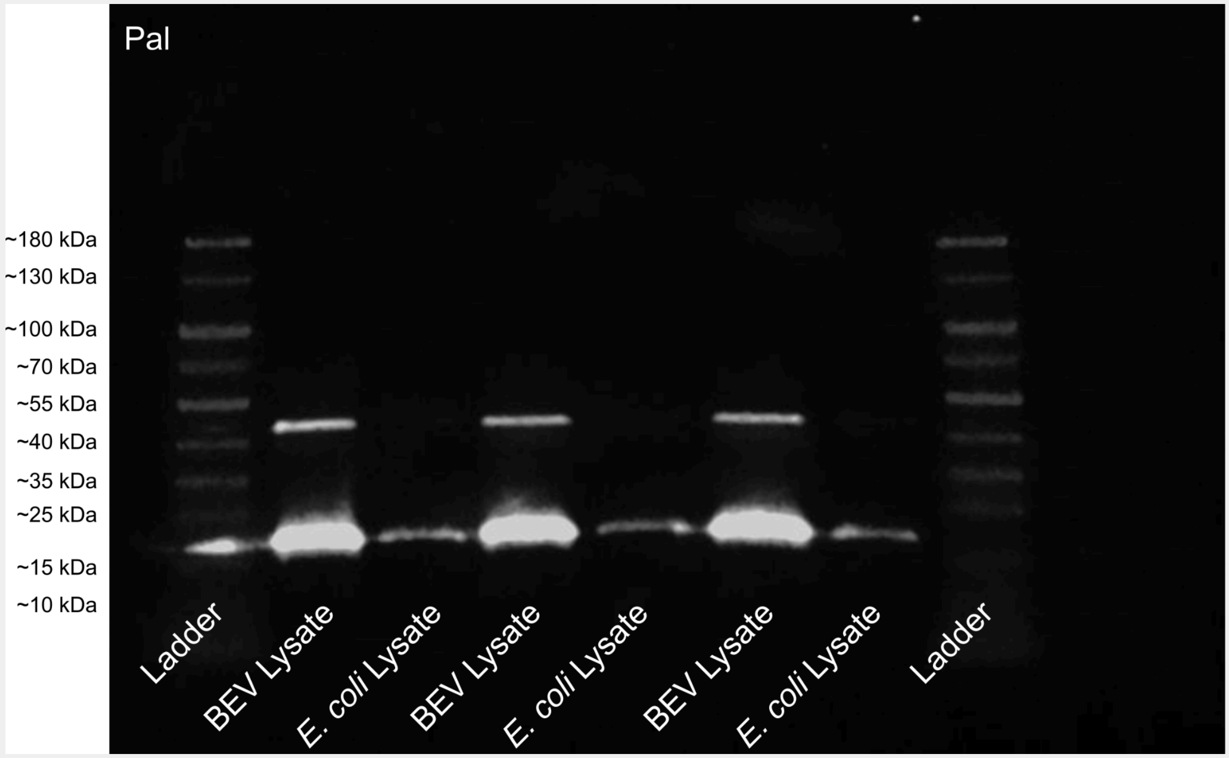


**Fig. S2: Full original fluorescent western blot image.** 1 µg of total protein from BEV lysates or *E. coli* whole cell lysates was added to each lane of the gel. Peptidoglycan-associated lipoprotein (PAL) was labeled using anti-Pal antisera from mice and an IRDye® 800CW goat anti-mouse IgG1-specific secondary antibody prior to obtaining the fluorescent image.

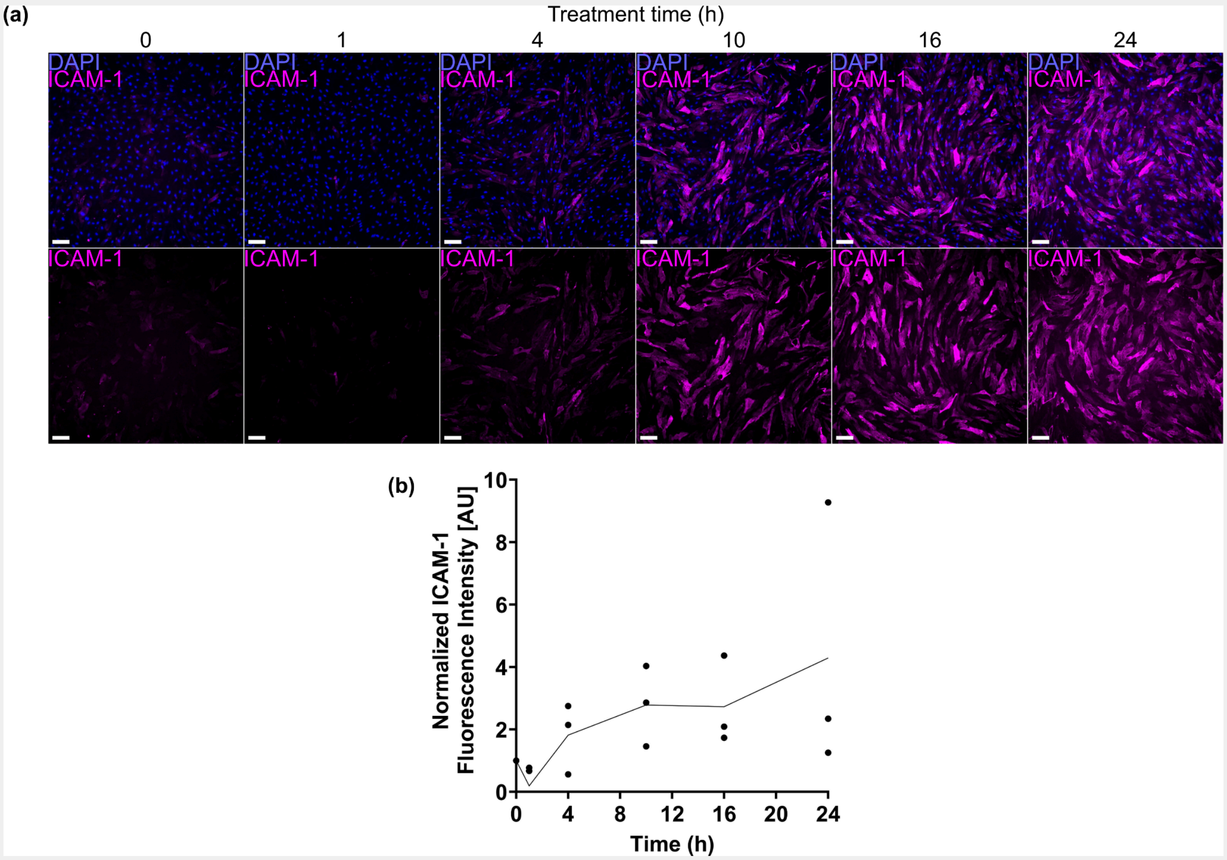


**Fig. S3: BMEC expression of ICAM-1 increases over the initial 10 h after LPS exposure and remains elevated for up to 24 h.** (**a**) Immunofluorescence images of BMECs seeded in a multiwell plate following treatment with 100 ng $ml^{-1}$ LPS for different periods of time. Cells were stained for nuclei (blue) and ICAM-1 (magenta). Scale bars = 100 µm. (**b**) Plot of fluorescence intensity of ICAM-1 divided by cell count and normalized to the 0 h time point.

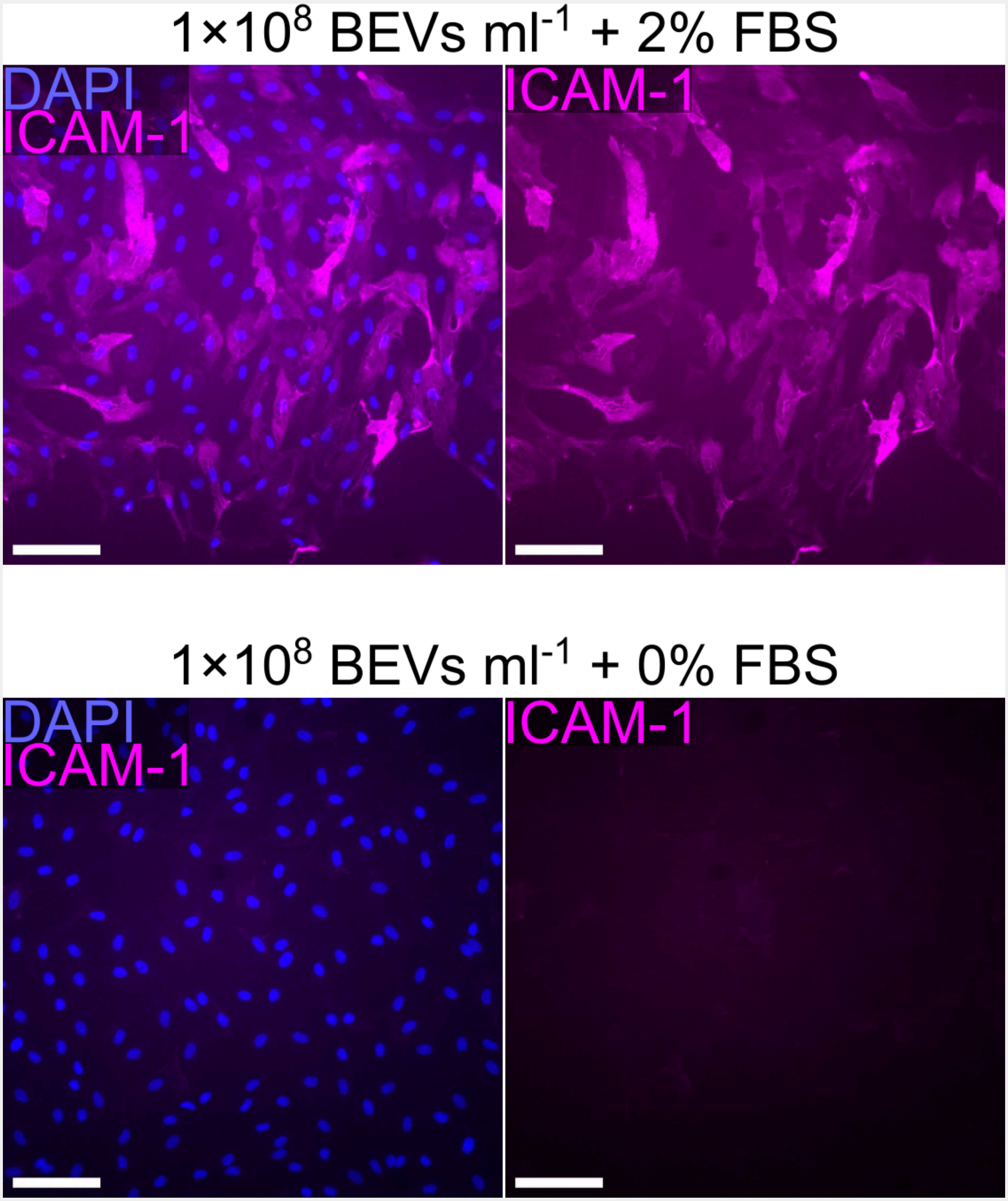


**Fig. S4: BMECs require a serum component in order to sense BEVs.** Immunofluorescence images of BMECs seeded in a multiwell plate following 16–17 h treatment with *E. coli*-derived BEVs at a concentration of $1\times10^{8}$ particles $ml^{-1}$. Cells were stained for nuclei (blue) and ICAM-1 (magenta). Culture medium was supplemented with 2% FBS (top) or without serum (bottom). Scale bars = 100 µm.

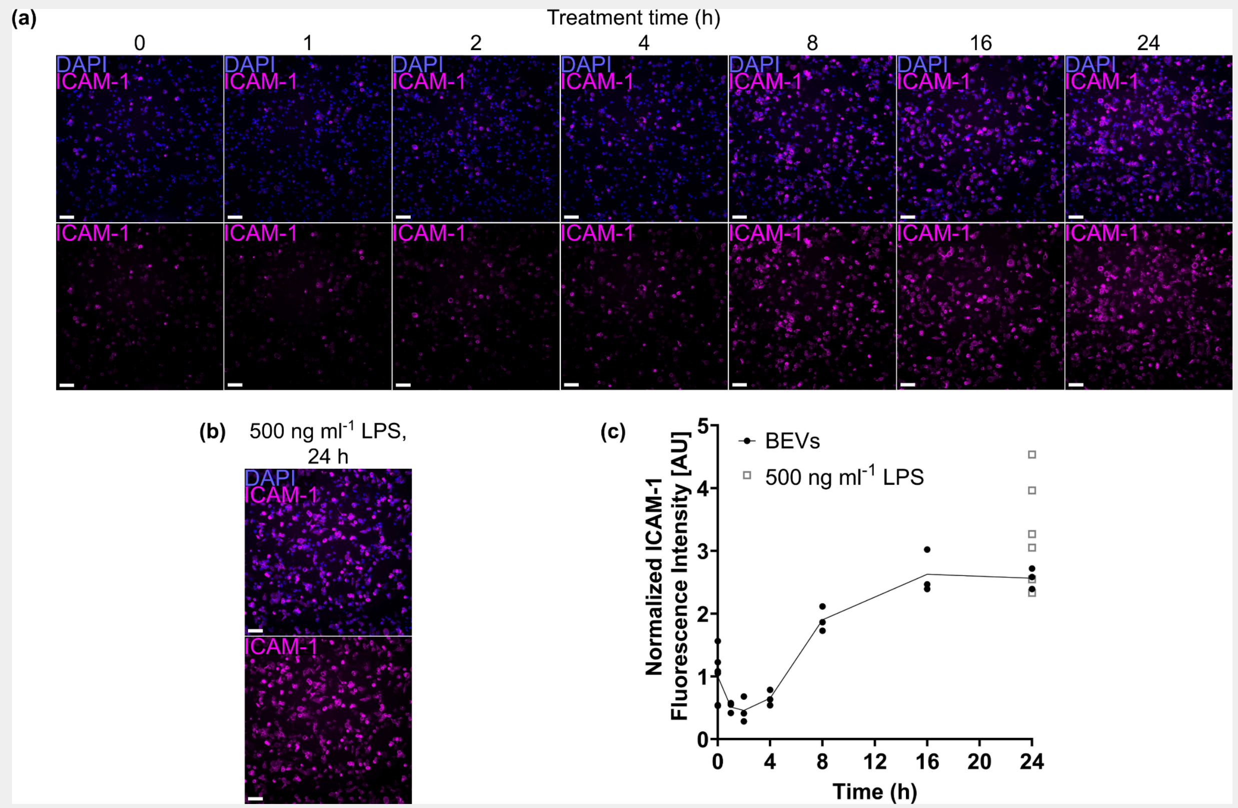


**Fig. S5: THP-1 macrophage expression of ICAM-1 increases over the initial 8 h after BEV exposure and remains elevated for up to 24 h.** (**a**) Immunofluorescence images of macrophage-like THP-1s seeded in a multiwell plate following treatment with $1\times10^{8}$ BEVs $ml^{-1}$ for different periods of time. Cells were stained for nuclei (blue) and ICAM-1 (magenta). (**b**) Immunofluorescence images of macrophage-like THP-1s seeded in a multiwell plate following 24 h treatment with 500 ng $ml^{-1}$ LPS, which was used as a positive control for ICAM-1 upregulation. Staining is identical to panel (a). (**c**) Plot of fluorescence intensity of ICAM-1 divided by cell count and normalized to the 0 h time point. Data points from the 24 h treatment with 500 ng $ml^{-1}$ LPS were included as a positive control. Scale bars = 100 μm.

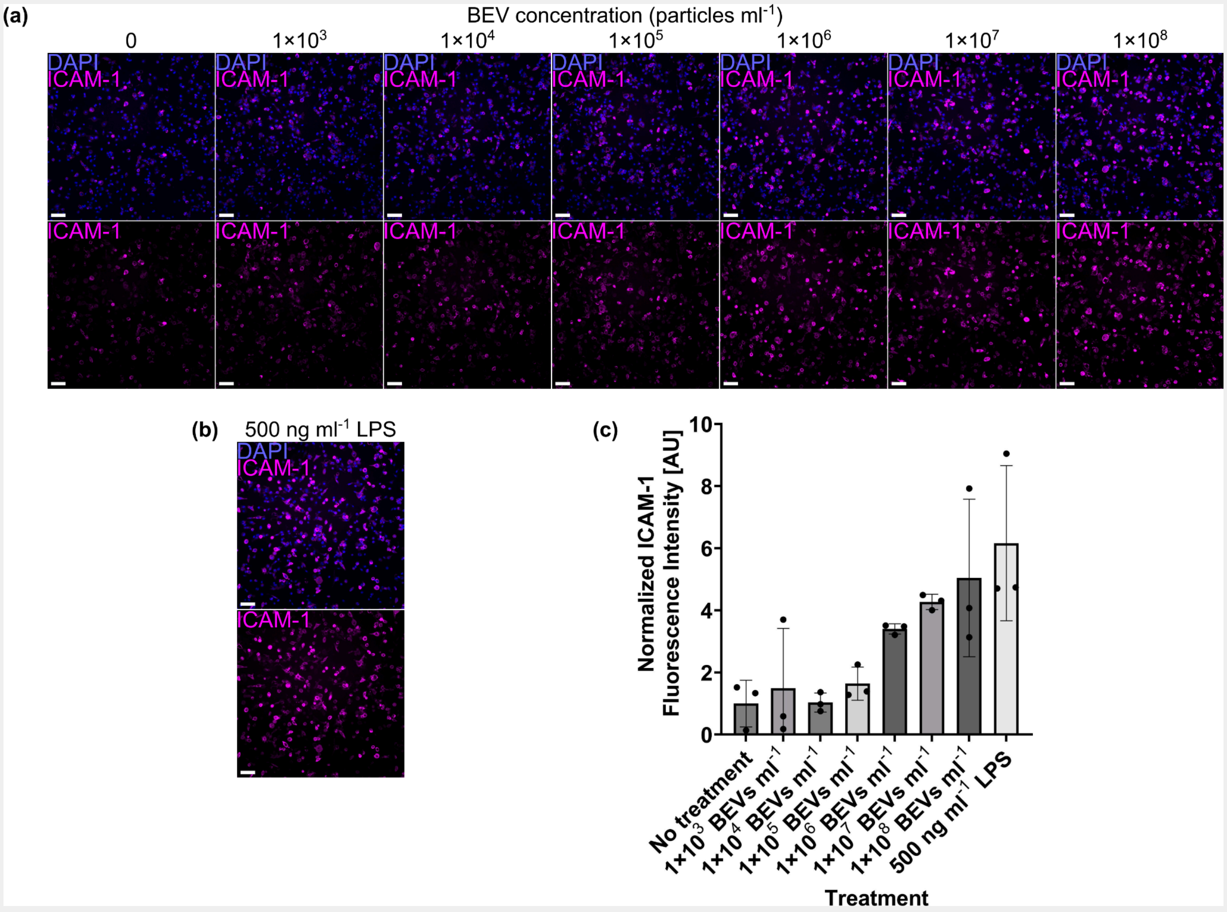


**Fig. S6: THP-1 macrophage expression of ICAM-1 increases due to BEV treatment in a dose-dependent manner** (**a**) Immunofluorescence images of macrophage-like THP-1s seeded in a multiwell plate following 16 h treatment with increasing concentrations of BEVs. Cells were stained for nuclei (blue) and ICAM-1 (magenta). (**b**) Immunofluorescence images of macrophage-like THP-1s seeded in a multiwell plate following 16 h treatment with 500 ng $ml^{-1}$ LPS, which was used as a positive control for ICAM-1 upregulation. Staining is identical to panel (a). (**c**) Barplot of fluorescence intensity of ICAM-1 divided by cell count and normalized to the “no treatment” condition. Scale bars = 100 μm. Error bars = s.d.

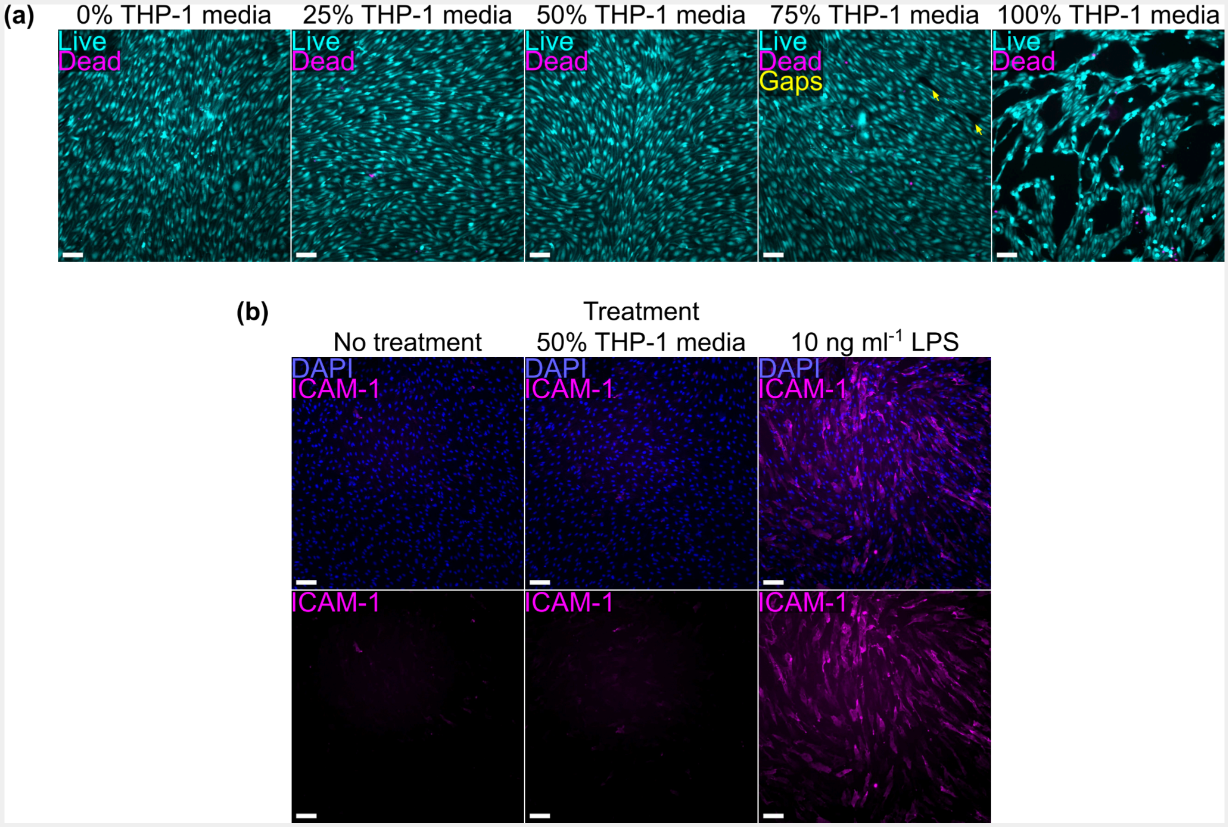


**Fig. S7: BMECs tolerate mixtures consisting of 50% THP-1-conditioned media.** (**a**) Immunofluorescence images of BMECs seeded in a multiwell plate following 16–18 h treatment with different mixtures of hECSR medium and conditioned medium derived from macrophage-like THP-1s. Live cells were labeled with calcein AM (cyan) and dead cells were labeled with ethidium homodimer-1 (magenta). Small gaps in the endothelial monolayer are labeled with arrows (yellow). (**b**) Immunofluorescence images of BMECs seeded in a multiwell plate following 16 h treatment with hECSR medium or 50% conditioned medium derived from macrophage-like THP-1s. Cells were stained for nuclei (blue) and ICAM-1 (magenta). 10 ng ml$^{-1}$ LPS treatment was used as a positive control for ICAM-1 upregulation. Scale bars = 100 µm.
